# Aggregation controlled by condensate rheology

**DOI:** 10.1101/2021.11.05.467474

**Authors:** Wolfram Pönisch, Thomas C.T. Michaels, Christoph A. Weber

## Abstract

Biomolecular condensates in living cells can exhibit a complex rheology including viscoelastic and glassy behaviour. This rheological behavior of condensates was suggested to regulate polymerisation of cytoskeletal filaments and aggregation of amyloid fibrils. Here, we theoretically investigate how the rheological properties of condensates can control the formation of linear aggregates. To this end, we propose a kinetic theory for linear aggregation in coexisting phases, which accounts for the aggregate size distribution and the exchange of aggregates between inside and outside of condensates. The rheology of condensates is accounted in our model via aggregate mobilities that depend on aggregate size. We show that condensate rheology determines whether aggregates of all sizes or dominantly small aggregates are exchanged between condensate inside and outside on the time-scale of aggregation. As a result, the ratio of aggregate numbers inside to outside of condensates differs significantly. Strikingly, we also find that weak variations in the rheological properties of condensates can lead to a switch-like change of the number of aggregates. These results suggest a possible physical mechanism for how living cells could control linear aggregation in a switch-like fashion through variations in condensate rheology.

**SIGNIFICANCE:** The intracellular space can be organized through phase-separated condensates that often exhibit rheological properties reminiscent of complex fluids. These condensates can affect biochemical processes such as the formation of linear aggregates, in particular biofilaments or amyloids. Here, we propose a theoretical model for how condensate rheology can control the irreversible formation of linear aggregates. A key finding is that size and number of aggregates change in a switch-like fashion upon weak changes in condensate rheology. Our model paves the way to unravel the physiochemical mechanisms of how the rheology of condensates can control aberrant protein aggregation. Such mechanisms may explain how rheological changes, such as ageing or the transition to dormancy, give rise to diseases related to protein aggregation.

## INTRODUCTION

The formation of linear aggregates plays an important role in many biological processes. Examples are biofilm formation (1, 2), the assembly of cytoskeletal filaments (3–5), and amyloids (6). The latter process is involved in a wide range of common and currently incurable diseases, such as Alzheimer’s, Parkinson’s, and amyloidosis (7–9).

Various theoretical models were proposed that capture key steps of the aggregation kinetics in vitro. An example is the pioneering model of Oosawa and Asakura for polymerization. In this model, linear aggregates can form via primary nucleation and grow at their ends. This model was successfully applied to actin and tubulin polymerization (3, 10). Ferrone and Eaton extended this model by introducing secondary pathways and applied to sickle hemoglobin polymerization (11, 12). More recently, related models were applied to describe amyloid formation (13–17). All such models capture the kinetics of aggregate in homogeneous environments.

However, many of such aggregation processes occur in living cells and living cells are strongly heterogeneous environments. The heterogeneity of intra-cellular space is for example due to condensates that form by phase separation (18–20). Such condensates can for example emerge as a response to cellular stress (21–23) and play a vital role during many biochemical processes. Examples are the enrichment of proteins (24, 25), the acceleration of gene expression (26) and enhanced drug resistance (27). Recent experimental evidence suggests that condensates also influence the formation of aggregates. For example, phase-separated compartments affect the self-assembly of DNA nanotubes (28) and can initiate or inhibit the formation of cytoskeletal filaments (24, 29–31), and amyloid fibrils (32).

In a recent study, a theoretical model was proposed for the irreversible aggregation kinetics inside and outside of a condensate (33). This model restricts to the case where exclusively monomers are exchanged between the condensate and its environment, but neglects aggregate diffusion. Though aggregates diffusive more slowly than small monomers in liquid phases due to their larger viscous drag, aggregates get still exchanged through the phase boundary of the condensate. However, many protein condensates exhibit complex rheological properties such as viscoelastic behavior or even glass-like ageing (34, 35). As a result of such complex rheology, the exchange of big aggregates can be significantly suppressed on the time-scale of aggregation and thereby alter the aggregation kinetics. It remained unclear to which degree the aggregation kinetics is affected by condensate rheology and what happens if the rheological properties change.

In our study, we investigate how the exchange of both monomers and aggregates between the two coexisting phases affects the kinetics of linear aggregation and how this exchange depends on condensate rheology. The rheological properties of condensates are accounted for by an aggregate mobility *ξ*_*i*_ that depends on aggregate size *i* (see Eq. (12)). Using our model, we find two distinct regimes with qualitatively different behaviours for the size distribution of linear aggregates. The regimes depend on whether all aggregates or only small aggregates are exchanged faster than the aggregation time-scale. Another key factor is the partitioning between the phases which is determined whether aggregates either interact over their complete length with the phase-separating material or exclusively with their ends. In our work, we develop the corresponding thermodynamic and kinetic theory to study how condensate rheology can affect irreversible formation of linear aggregates that interact differently with the phase-separating material. We report differences in size distributions and aggregate mass ratio between the two phases and reveal a switch-like change of the aggregation kinetic upon weak changes in condensate rheology.

## THEORETICAL MODEL FOR RHEOLOGY-DEPENDENT AGGREGATION

In the following, we introduce a set of master equations to describe linear aggregation in the presence of a phase-separated condensate with various rheological properties. We also derive the governing relationships for how monomers and aggregates partition and are transported between the inside and outside of the condensate. The rate of aggregate exchange between the phases is considered to have a specific dependence on aggregate size which is a result of a specific condensate rheology. Thus, the resulting model allows us to discuss how rheological properties of condensates can affect the kinetics if aggregation.

### Master equation of irreversible linear aggregation

To describe the temporal evolution of irreversible linear aggregation, we introduce a set of master equations for the aggregate concentration 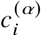 (units of molar concentration M). Here, *i* indicates the aggregate size, while *α* denotes the inside (*α* = I) and outside (*α* = II) of the condensate, also called phase I and phase II. The evolution of monomer and aggregate concentrations in each phase is given by

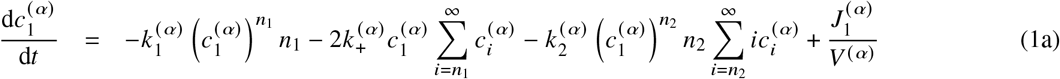

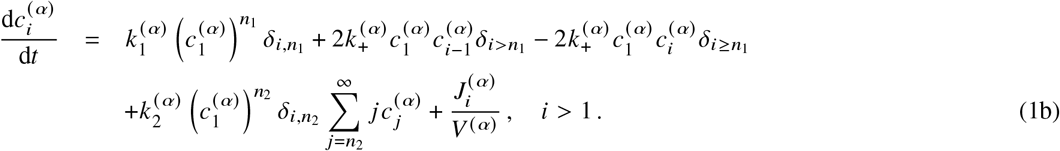

The terms including the rate constant 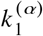 correspond to primary nucleation, leading to the initial formation of aggregates. During primary nucleation, *n*_1_ monomers assemble one aggregate of size *n*_1_, where *n*_1_ is called the reaction order. Monomers can also bind to the ends of aggregates with a rate constant 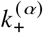, creating aggregates with a size *i* > *n*_1_. Additionally, the subunits of aggregates of size *i* ≤ *n*_2_ act as nucleation sites for new aggregates of size *n*_2_ to form by a process called secondary nucleation occurring with a rate constant 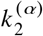. In general, the rate constants 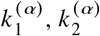 and 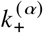 can be different between both phases I and II. For example, if reactions between monomers and aggregates are limited by their diffusion, the rates might depend on the rheology of the two phases. For simplicity, we consider phase-independent rate constants and write *k*_1_, *k*_2_ and *k*_+_. In Fig. 1 *A*, we provide a graphical representation of primary and secondary nucleation and aggregate elongation.

**Figure 1:**
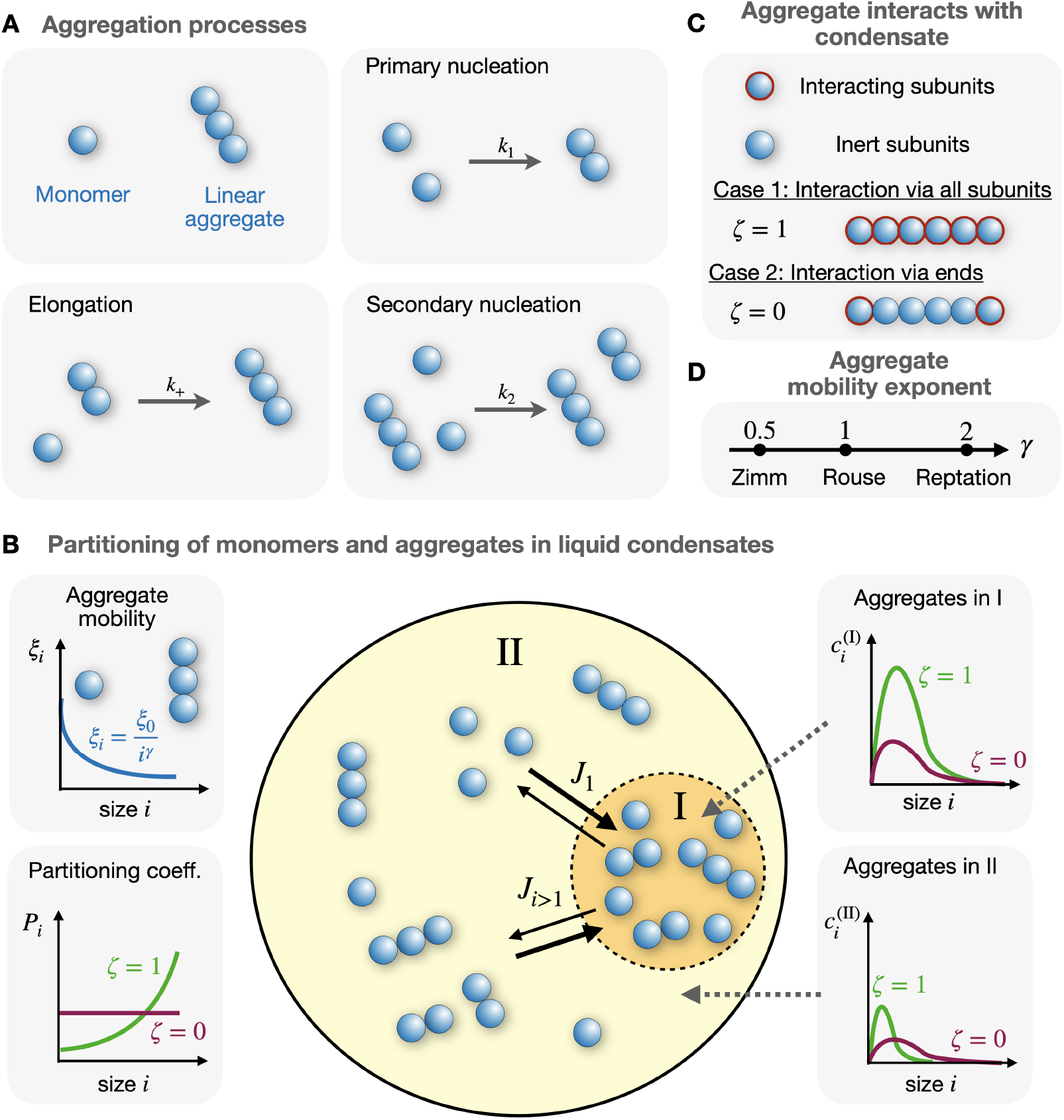
Illustration of our model for linear aggregation in the presence of a phase-separated condensates with varying rheology. (A) Elementary processes driving the formation of monomers to linear aggregates: primary nucleation with rate constant *k*_1_, elongation with rate constant *k* and secondary nucleation with rate constant *k*_2_. (B) Monomers and linear aggregates within phase-separated condensates. Monomers and aggregates of size *i* are exchanged between phase I to phase II by diffusion with a rate *j*_*i*_ [units: 1/s]. The aggregate size dependence of the aggregate mobility *ξ*_*i*_ (Eq. (12)) and the partition coefficient *P*_*i*_ (Eq. (4)) control the aggregation kinetics and thus the profile of the aggregate concentrations 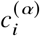 inside and outside of the condensate, *α* = I, II. Both also depend on how linear aggregates interact with the phase-separating components A and B, (C) either all aggregate subunits interact with the condensate (binding parameter *ζ* = 1) or only aggregate ends interact with the condensates (*ζ* = 0. (D) Sketch of the different regimes of the mobility exponent *yγ* of the linear aggregates, dependent on the aggregate concentration and hydrodynamic interactions.

The master equations Eqs. (1) are motivated by previous studies of cytoskeletal polymerisation (3, 10) and amyloid fibril aggregation (14, 15, 17, 36, 37). Eq. (1a) governs the temporal dynamics of monomers and Eq. (1b) describes the temporal dynamics of the aggregates of length *i >* 1. Phase coexistence dictates the use of the chemical activities 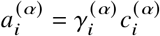 of the aggregates in the reaction steps, where 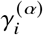 denotes the chemical activity coefficient. This accounts for the coupling between aggregate partitioning and nucleation and growth of aggregates (38). However, in Appendix D, we show that both processes can decouple for irreversible processes leading to Eqs. (1).

There is a diffusive total exchange rate *J*_*i*_ that describes the exchange of aggregates of size *i* between the interior and exterior of the condensate (see Fig. 1 *B*). The total exchange rate conserves monomer and aggregate mass as well as particle numbers, and thus obeys 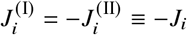. Here, we consider the case of dilute aggregates which implies that the exchange of aggregates between the inside and outside does not affect the condensate volumes *V* ^(I)^ = *V* − *V* ^(II)^, where *V* is the system volume.

The stationary state of Eq. (1) is a non-equilibrium steady state. In our work, we explicitly focus on aggregation kinetics where the time-scale of reaching thermodynamic equilibrium exceed the time-scale of interest where most monomers are depleted. A well-studied example for the latter case is the aggregation of amyloid fibrils. Motivated by this class of aggregation processes, we consider a special case where the reaction kinetics for nucleation and growth in each phase (i.e., 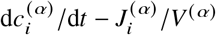) have the same mathematical form as in a homogeneous system and are independent of partitioning of monomers and aggregates.

We also introduce the aggregate number concentration, defined by

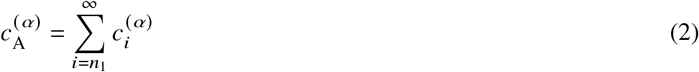

and the aggregate mass concentration

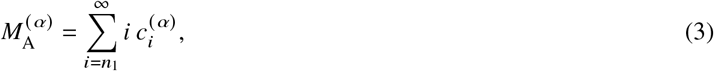

which correspond to the zeroth and first moment of the aggregate concentration 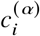, respectively. Details of the numerical solution of the master equations are given in Appendix A.

The interplay of linear aggregation and the flux of aggregates between the inside and outside of the condensate controls the aggregate concentration profile within the two phases I and II. This interplay depends on the partitioning of monomers and aggregates as well as on the rate of diffusive exchange. In the following, we derive an expression for the partition coefficient of aggregates as a function of their size, and then derive the flux law for monomer and aggregate exchange.

### Phase separation and partitioning of linear aggregates

We define the partition coefficient at phase equilibrium as

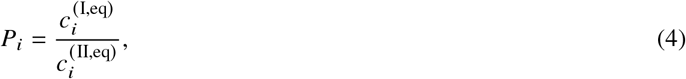

of aggregates of size *i* and concentration 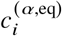 are the equilibrium concentrations inside (*aα* = I) or outside (*α* = II) of the condensate. At phase equilibrium, chemical potentials, *µ*_*i*_ ≡ *i ν∂ f* /*∂ ϕ* _*i*_ (*v* is the molecular volume of a monomer), are equal between both phases:

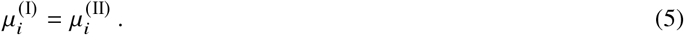

To determine the chemical potential, we consider an incompressible mixture composed of two phase-separating components *A* and *B*, and aggregating monomers. For example, phase I is A-rich (B-poor), while phase II is B-rich (A-poor). The monomers are prone to undergo irreversible aggregation into linear aggregates of size *i*. We study the case where the monomers and the resulting aggregates are highly diluted with respect to components *A* and *B*, which is consistent with physiological conditions, e.g. for amyloid-*β* monomers and fibrils (see Appendix B). To derive the partitioning of the aggregates, we use a free energy that is qualitatively similar to the Flory-Huggins free energy (39, 40), but does not rely on mean field approximations of the highly dilute linear aggregates (41). The derivation of the free energy of dilute aggregates of size *i* is provided in Appendix C.

This free energy density is given by

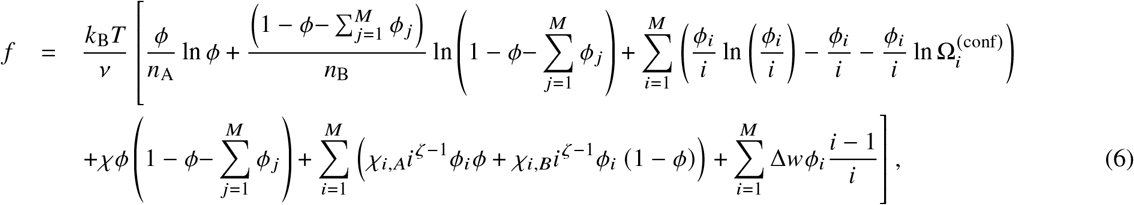

where *ϕ*_A_ = *ϕ*and 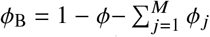 denote the volume fractions of components *A* and *B*, respectively, and *n*_A_ and *n*_B_ are the sizes of components *A* and *B* measured in terms of the molecular volume of monomers denoted as *v*. Moreover, *ϕ*_*i*_ is the volume fraction of the aggregates of size *i* and *M* is the maximal aggregates size, i.e. 1 ≤ *i* ≤ *M*. Note that *ϕ*_*i*_ « 1 since the linear aggregates are dilute.

The first two terms in Eq. (6) correspond to the mixing entropy of the components *A* and *B*. The sum over the aggregate size *i* involves contributions from the mixing entropy of the aggregates and entropic contributions related to the number of possible aggregate configurations, 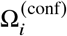, which depends on the size *i*. The terms including the parameters *χ, χ*_*i,A*_ and *χ*_*i,B*_ represent enthalpic contributions due to the interactions between the components *A, B* and the aggregates of size *i* with the respective interaction parameters *χ,χ*_*i,A*_ and *χ*_*i,B*_. Since the linear aggregates are considered to be dilute, we have neglected aggregate-aggregate interactions in the free energy. In some experimental studies on the molecular weight dependence of the interaction parameter in polymer-polymer-good-solvent systems it has been suggested that the interaction parameters depend on polymer length by the equations 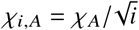and 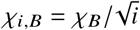 (42). For simplicity, we will neglect any aggregate size dependence of the interaction parameters, setting *χ*_*i,A*_ = *χ*,_*A*_ and *χ*,_*i,B*_ = *χ*,_*B*_, as suggested in experimental measurements in polymer-liquid crystal mixtures (43), but we note that our model can be extended in a straightforward way to account for different size dependencies of *χ*,_*i,A*_ and *χ*_*i,B*_.

We also introduce the parameter *ζ*, which we refer to as binding parameter in the following. The binding parameter *ζ* characterizes how the aggregate subunits are interacting with the phase-separating components *A* and *B*: *ζ* = 1 corresponds to the case where all subunits can bind to *A* and *B* molecules, while *ζ* = 0 is the case where only the subunits at the linear aggregate ends can bind to *A* and *B* molecules (see Fig. 1 *C*). This could play a role for subunits with hydrophobic binding sites that are buried when the subunit is within the bulk of a linear aggregate, but freely available for binding to *A* and *B* for subunits at the aggregate ends. The internal free energy Δ*w* describes the free energy of each bond in a polymer of length *i*. Since both coexisting phases are liquids, Δ*w* is phase-independent.

The chemical potential of aggregates of size *i* can be calculated using Eq. (6), leading to

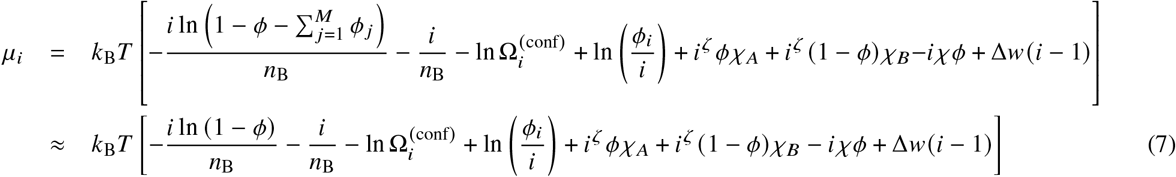

At equilibrium, the difference of chemical potentials between two separated phases, (I) and (II) (see Fig. 1 *B*)

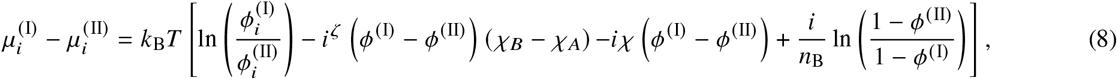

is zero. Using this condition, we find for the partition coefficient

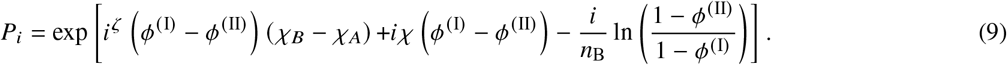

At phase equilibrium, it fulfils

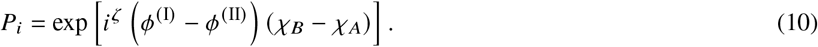

Since we assume that the aggregate concentration 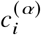 is proportional to the aggregate volume fraction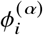, Eq. (10) implies 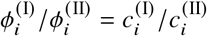. For the binding parameter *ζ* = 1, the partition coefficient increases monotonically with aggregate size, assuming that (*ϕ*^(I)^ − *ϕ*^(II)^) (*χ*_*B*_ − *χ*_*A*_) > 0, and hence, *P*_*i*_ increases with the molecular weight of aggregates. This increase is in accordance with previously reported experimental studies on hydrophobic compounds in polymer-water mixtures (44) and dilute polymers between cylindrical pores and an exterior solution (45). While in both studies, the partition coefficient increases exponentially with molecular weights, this is only true for a small range of molecular weights. We also consider the case ζ = 0, corresponding to the limit in which *A* and *B* exclusively bind to the aggregate ends (see Fig. 1 *C*) and as a result, the partition coefficient is independent of aggregate size *i*.

### Flux of linear aggregates between the inside and outside of the condensate

Near equilibrium, the total flux of aggregates of size *i* is proportional to the difference of the corresponding chemical potentials inside and outside the condensate and can be written as (33):

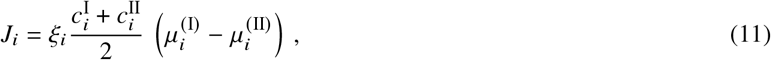

where the dependence on 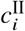 ensures that the mobility *ξ* _*i*_ is constant in the dilute limit. This mobility characterizes how fast a phase of size *V* ^I 1/3^ is homogenized suggesting a phenomenological relationship to the diffusion coefficient of the form *D*_*i*_ ∼ *ξ*_*i*_ (*V*^I^) ^1/3^ if the transport of aggregates inside is the rate limiting step.

To account for the impact of condensate rheology on aggregation kinetics, we introduce a mobility that depends on aggregate size *i*. In the following, we consider a power law dependence of the form:

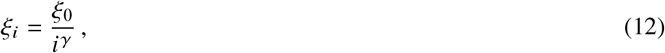

where *ξ*_0_ is the mobility prefactor and *γ* denotes the exponent of the power-law (see Fig. 1 *B* and Fig. 1 *D*). In our work, we distinguish between three fundamentally different rheological behaviors corresponding to different interactions among the polymeric components (Figure 1 *D*). For polymers interacting via hydrodynamic interactions, the exponent is *γ* = 0.5, which corresponds to the Zimm regime. In the absence of hydrodynamic interactions, solely accounting for single polymer friction with the solvent, *γ* = 1, which is refereed to to the Rouse regime. The case *γ* = 2 describes polymers that are entangled by neighboring polymers which can escape via reptation (46, 47). We also consider non-integer valued rheology exponent exponent *y* to account for complex polymer melts or micro phase-separated mixtures.

Based on the mobility *ξ*_*i*_, we can derive a characteristic time-scale for the exchange of aggregates between the condensate inside and outside (see Appendix F):

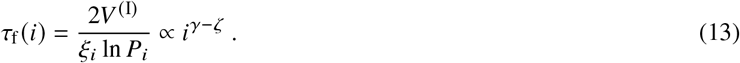

When the binding parameter *ζ* is equal to the mobility exponent *γ* i.e. *γ* = *ζ*, the characteristic time of aggregate exchange is independent of aggregate size. For *γ > ζ* larger aggregates are exchanged slower between the condensate inside and outside than small ones, while the opposite holds for *γ < ζ*.

The characteristic exchange of aggregates *τ*_f_ *i* allows us to enforce phase equilibrium depending on aggregate size *i*. In particular, we consider two limits: on short time-scales *t < τ*_f_ *i*, aggregates of size *i* are not exchanged between the phases. In this case, we set the exchange total exchange rate *J* _*i*_ = 0 implying that aggregates nucleate and grow in each phase independently of the other phase. In contrast, on large time-scales *t ≥ τ*_f_ (*i)*, aggregates are at phase equilibrium and thus, at each time point, the relative concentration of aggregates follow the partition coefficient (see Eqs. (4) and (10)).

## RESULTS AND DISCUSSION

In the following, we investigate the aggregation dynamics of dilute linear aggregates in a system where two phases coexist and study how we can control the outcome of aggregation by varying the condensate rheology via changes of the mobility prefactor *ξ*_0_ and exponent γ. To this aim, we solve the master equations (see Eqs. (1)) numerically; see Appendix A for details. Our theory can describe the irreversible nucleation and growth kinetics of linear aggregates in the presence of phase-separated condensates. In the following, we choose parameters consistent with experimental measurements on A*β*42 monomers forming amyloid fibrils (details see Appendix A). For droplets of sizes around 1 − 10 *µ*m, we find that the mobility prefactor *ξ*_0_ in Eq. (11) has values around *ξ*_0_≈10^3^− 10^4^ *µ*m^3^ s (see Appendix E). For the initial monomer concentration, we pick values that are comparable to in-vitro studies on amyloid kinetics, i.e., 4 *µ*M (32). This concentration is far above estimates for in-vivo concentrations which are around 10 - 200 pM (48, 49). The choice of a *µ*M-ranged monomer concentration enables us to scrutinize our results by in-vitro experiments where typical aggregation times are in order of hours. Additionally, we consider condensates that are small compared to the system size, *V* ^(I)^ *V* ≃10−^2^« 1, to highlight that even small volumes can have a significant impact on aggregation.

### Condensate rheology controls aggregation kinetics and distribution

To identify how condensate rheology affects monomer and aggregate partitioning, we solve the master equations (see Eq. (1a) and Eq. (1b)) and quantify the main features of the linear aggregate size distribution as a function of the mobility prefactor *ξ*_0_ and exponent *γ* (see Eq. (12), Fig. 2 and Fig. S1).

**Figure 2:**
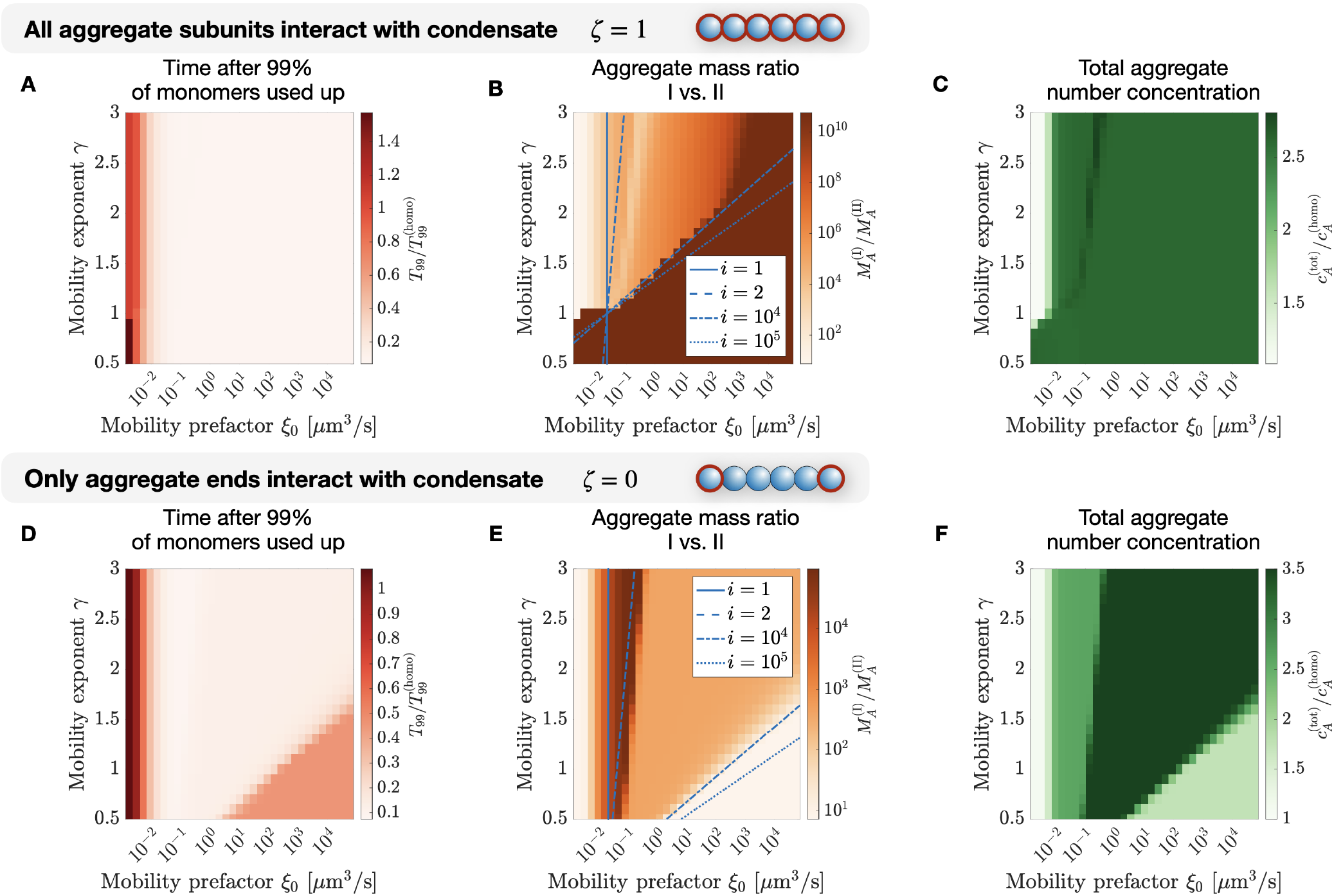
Diagrams of how condensate rheology affects linear aggregate distribution. For the binding parameter *ζ* = 1 (the partition coefficient is given by *P*_*i*_ = exp 2*i*) and *ζ* = 0 (with *P*_*i*_ = exp 2), the master equations (see Eq. (1a) and Eq. (1b)) is solved and (A,D) the aggregation time *T*_99_, (B,E) aggregate mass ratio and (C,F) the total aggregate number concentration 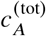 are given after the time *T T*_99_, corresponding to the time it takes 99 % of monomers in the system to assemble to linear aggregates. The aggregation time *T*_99_ is normalised by the time ^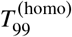^ it takes to for the monomers to aggregate in the homogeneous case with only a single phase, but the identical initial total monomer concentration. Analogously, the total aggregate number concentration is normalised by the total aggregate number concentration of the homogeneous case. The quantities are investigated for different values of the mobility exponent *γ* and prefactor *ξ* _0_ (see Eq. (12)). The blue curves in show when the aggregation time ^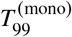^ in the case of exclusive and instantaneous monomer exchange (see supporting information S1.2) and the aggregate exchange time (see Eq. (13)) are the same as a function of aggregate sizes *i*. Here the total initial monomer concentration is 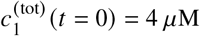 (see Eq. (14)) and the volumes are *V* ^(I)^ = 10.1 *µ*m^3^ and *V* ^(II)^ = 1000 *µ*m^3^.

Initially, we focus on the case *ζ* = 1, which corresponds to aggregates binding to the phase-separating components *A* and *B* with all their subunits. Setting the binding parameter *ζ* = 1 in Eqs. (10) makes the partition coefficient *P*_*i*_ size-dependent. We assume that *P*_*i*_ *>* 1, so that monomers and aggregates are accumulated inside the condensate (phase I). We start our investigation by initializing the system exclusively with monomers at their equilibrium concentration inside and outside the condensate, fulfilling 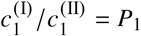. We then allow the monomers to form aggregates and both monomers and aggregates are exchanged between the inside and outside of the condensate if their exchange time *rτ*_f_ *> t* (see Eq. (13)) with the time *t*. The reduction of the monomer concentration leads to monomer and aggregate fluxes between the two phases to fulfil the partition coefficient. We numerically solve the master equations until 99 % of the initial total monomer concentration 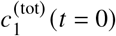 in the system have assembled into aggregates of size *j >* 1. This threshold allows us to define the point at which most monomers in the system have assembled into aggregates and the remaining dynamics of the system are dominated by monomer and aggregate exchange between the two phases. The corresponding time _99_ is determined by the condition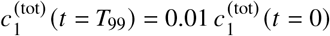. Here, the total monomer concentration is given by

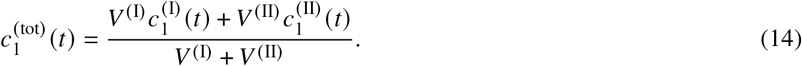

Since we do not allow for any fragmentation or aggregation processes that are not involving monomers, the aggregation process slows down the fewer monomers are left and stops when no more monomers are available. In Fig. 2 *A*, we show the time _99_ and we find that this time is dominantly dependent on the mobility prefactor *ξ*_0_ for *ζ* = 1. For small values of *ξ*_0_, the time approaches the value of the homogeneous system with a single phase only. For small mobility prefactor *ξ*_0_ and small mobility exponent *γ*, we find that the time can even exceed the aggregation time in the homogeneous case. For higher values of *ξ*_0_, the time *T*_99_ decreases, implying that condensates accelerate aggregation due to the accumulation of monomers and linear aggregates within phase I, as previously described (33). For *ξ*_0_ *>* 0.01 *µ*m^3^ s, the time _99_ is not dependent on the mobility exponent *γ*. This result suggests that the time *T*_99_ is dominantly influenced by the monomer exchange, which is independent of *γ*, and not by the aggregate exchange, which is controlled by the mobility exponent *γ*.

We next investigated the distribution of aggregate mass in the two phases by looking at the ratio of the aggregate mass concentrations (see Eq. (3) and Fig. 2 *B*) and the aggregate mass concentration outside of the condensate in phase II (see Fig. S1 *A*). We find that independently of the mobility and since *P*_*i*_ *>* 1, the mass ratio is higher than 1, hence aggregates are accumulated within the condensate (phase I). Additionally, we find two regimes: In the first regime, given by small values of *γ* and large enough values of *ξ*_0_, the mass ratio is very high and exceeds 10^10^ and the aggregate mass concentration outside of the condensate is around 10^−8^ times smaller than the concentration for the single phase case. This corresponds to essentially all linear aggregates being located within the condensate. For higher values of *γ* and smaller values of *ξ*_0_, the mass ratio is in the order of 10^2 −^10^4^ and considerably smaller. We observe similarly pronounced differences in the ratio of mean aggregate size between the inside and outside of the condensate (see Fig. S1 *B*). We always observe that linear aggregates within the condensate (in phase I) have a larger mean size than aggregates outside of the condensate (in phase II), but for small enough *γ*, the size ratio is considerably higher than for larger *γ*.

The mass ratio and the mean aggregate size ratio are both quantities that have not yet equilibrated after 99 % of monomers have aggregated. The ratio of aggregates outside and inside of the condensate does not necessarily fulfil the partition coefficient *P*_*i*_ (see Eq. (10)) yet and the aggregate flux is then non-zero. Thus, we consider the total aggregate number concentration,

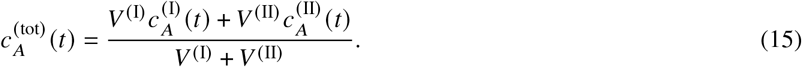

This quantity describes the total concentration of aggregates in the total volume, *V* ^(I)^ +*V* ^(II)^, and is near its equilibrium value at *t* = *T*_99_ since most monomers have assembled to aggregates and the irreversible aggregate production has almost ended. We show the total aggregate number concentration at time *T*_99_ in Fig. 2 *C*. We find that analogously to the time *T* _99_, the final total concentration 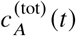 is only weakly dependent on the mobility exponent *γ*, but dependent on the mobility prefactor *ξ*_0_. For *ξ*_0_ *<* 0.01 *µ*m /s, we find that the aggregate number concentration is similar to the homogeneous case, except for *γ <* 1. Additionally, we find a weak non-monotonous dependence of the number concentration on *γ* for *ξ*_0_ *>* 0.1.

Until now, we only studied the case where all aggregate subunits interact with the condensate, corresponding to the binding parameter *ζ* = 1. We now consider the case *ζ* = 0, where aggregates interact exclusively with their ends with the condensate. The same quantities we studied before for *ζ* = 1, i.e. the aggregation time, the aggregate mass and mass ratio, the total aggregate concentration and the aggregate size ratio, are shown in Fig. 2 *D-F* and Fig. S1 *C-D*. For *γ >* 1.5, the time _99_ decreases with mobility prefactor *ξ*_0_ (see Fig. 2 *D*) and shows the same behaviour as for *ζ* = 1. For *γ <* 1.5, it initially decreases with *ξ*_0_, but then starts to increase again. The aggregate mass ratio (see Fig. 2 *E*) initially increases with *ξ*_0_, and decreases after reaching a maximum. It then stays constant with *ξ*_0_, before once more decreasing. As a result, we can identify four different regimes. The same regimes can also be found in the total aggregate number concentration Fig. 2 *F*, which shows a non-monotonous dependence on *ξ*_0_.

### Switch-like transition in the aggregation dynamics upon changes in condensate rheology

In the previous section, we showed that there is transition between different regimes in the aggregate mass ratio (see Fig. 2 *B,E*) as a function of the values of the mobility exponent *γ* and prefactor *ξ*_0_. In this section, we will show that this transition can occur in a switch-like fashion. To see this behavior, which is reminiscent of a non-equilibrium bifurcation, we first compare the time scales of aggregate exchange between with the condensate and of aggregation. We show that the location of the transition in Fig. 2 *B,E* depends on whether aggregation is quicker than the aggregate exchange. Next, we numerically solve the master equation near this transition and find that the transition is indeed switch-like. The switch-like behaviour is especially pronounced when all aggregate subunits interact with the condensate, *ζ* = 1.

We first consider the time 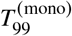 of aggregation. This is the time it takes to assemble 99 % of monomers in the case that only monomers are exchanged instantaneously between the inside and the outside of the condensate, while aggregates are not exchanged (see supporting material S1.2 for the analytical solution of this case). This corresponds to a high mobility exponent *γ* and mobility prefactor *ξ* _0_. When comparing the aggregation time to the aggregate size-dependent exchange time *τ* _f_ (see Eq. (13)), we find that only those aggregates with size *i* that fulfil 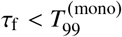 are transported fast compared to the timescale of aggregation. For a given *i*, this corresponds to

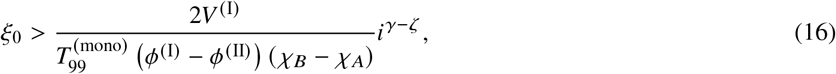

where we combined Eq. (10), Eq. (13) and supporting material S1.2.

### Properties of switch-like transition when all aggregate subunits interact with condensate (( = 1)

For *ζ* = 1, linear aggregates are dominantly transported from phase II to I, since the partition coefficient *P*_*i*_ increases exponentially with aggregate size *i* (see Eq. (10)). We now show that with the help of the relation given in Eq. (16), we can predict the location of the transition between the two regimes of the mass ratio in Fig. 2 *B*. Then, we show that the mass ratio changes dramatically over multiple orders of magnitude for small changes of the mobility exponent *γ*, which we interpret as a switch-like behaviour.

During the time of aggregation, for the most part, aggregates of size *i* that fulfil 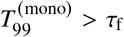 are transported between the two phases. In Fig. 2 *B*, we show blue curves that fulfil ^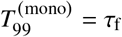^ for different values *i* and we find that for a certain range of size *i*, we can qualitatively predict where the transition between the two regions in Fig. 2 *B* happens. This implies that for small values of *γ* or high values of *ξ* _0_, aggregates of all sizes are exchanged between the inside and outside of the condensate, explaining the extremely high mass ratio in Fig. 2 *B*. For high values of *γ* or small *ξ* _0_, the relation in Eq. (16) is only fulfilled for small aggregate sizes *i*, hence only monomers and small linear aggregates are exchanged, reducing the mass ratio and affecting the aggregation within phase I. For small values of both the mobility coefficient *ξ* _0_ and the mobility coefficient *γ <* 1, monomers are exchanged slower than aggregates grow and as a result, the aggregation time is slower than in the homogeneous case (see Fig. 2 A). This difference is because aggregates that are generated in phase II are transported into phase I, while monomers remain in phase II. Within phase I, secondary nucleation and monomer pickup are slowed down since monomers are depleted. Eq. (16) allows us to derive a characteristic aggregate size for which material exchange happens faster than aggregation.

However, it is important to note that the mean aggregate size 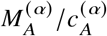 is not constant during aggregation, but time-dependent. To get an estimate of an characteristic aggregate size, we determine the maximal aggregate size. Therefore, we study the aggregate concentrations for the homogeneous aggregation (see supporting material S1.1 and Fig. S2) and find that the aggregate profile possesses a moving front. This front originates from the finite time the monomers have to aggregate. Aggregates with a size exceeding the front did not have sufficient time to assemble. The highest value of this front has the same order of magnitude as the aggregate size for which Eq. (16) best recovers the location of the transition in Fig. 2 *B*), *i* ≈ 10^4^− 10^5^. This observation suggests that the transition in the aggregate mass ratio (see Fig. 2 *B*) happens when all linear aggregates that exist in the system are exchanged faster between the condensate interior and exterior than the aggregation dynamics. We are now able to explain the two regimes (for *ξ* _0_ *>* 1 *µ*m^3^ s) found in Fig. 2 *B*, characterised by significant differences in the aggregate mass ratio. We conclude that in the region defined by small mobility prefactor *ξ* _0_ or large enough mobility exponent *γ*, monomers and small aggregates are dominantly exchanged between the two phases, since the time scale of aggregate exchange is too large compared to the time of aggregation for larger aggregates. Hence, aggregates remain outside of the condensate, decreasing the mass ratio. When the mobility exponent *γ* is reduced or the mobility prefactor *ξ* _0_ is increased, the time scale of aggregate exchange deceases until the largest aggregates can be exchanged on a time-scale faster than aggregation. This defines the/a second region for low values of *γ* or high *ξ* _0_ in the phase diagram which is characterised by a significantly increased aggregate mass ratio since all aggregates are transported into phase I.

For *ξ* _0_ *<* 1 *µ*m^3^ s, we also discover another transition that is linked to whether oligomers of size *i* = 2 are exchanged between the phases. For values of *ξ* _0_ that are smaller than the the transition marked by the blue dashed line in Fig. 2 B, oligomers are not exchanged while aggregation is happening.

We next investigate the behaviour in the vicinity of the transition in the region defined by *ξ* _0_ *>* 1 *µ*m^3^ s. In Fig. 3 *A*, we show the aggregate mass ratio as a function of the mobility exponent *γ*. The mass ratio shows a pronounced change with the mobility exponent *γ*, reminiscent of a switch between two different limits. The switch between both limits is reflected in the rate of change of the mass ratio as a function of the mobility exponent *γ* (see Fig. 3 *C*), possessing a pronounced peak. Additionally, the position *γ* _peak_ of the peak is increasing with the mobility prefactor *ξ* _0_. This suggests that, by varying its rheological features slightly, a cell can alter the qualitative outcome of the aggregate distribution near the region where the transition happens.

**Figure 3:**
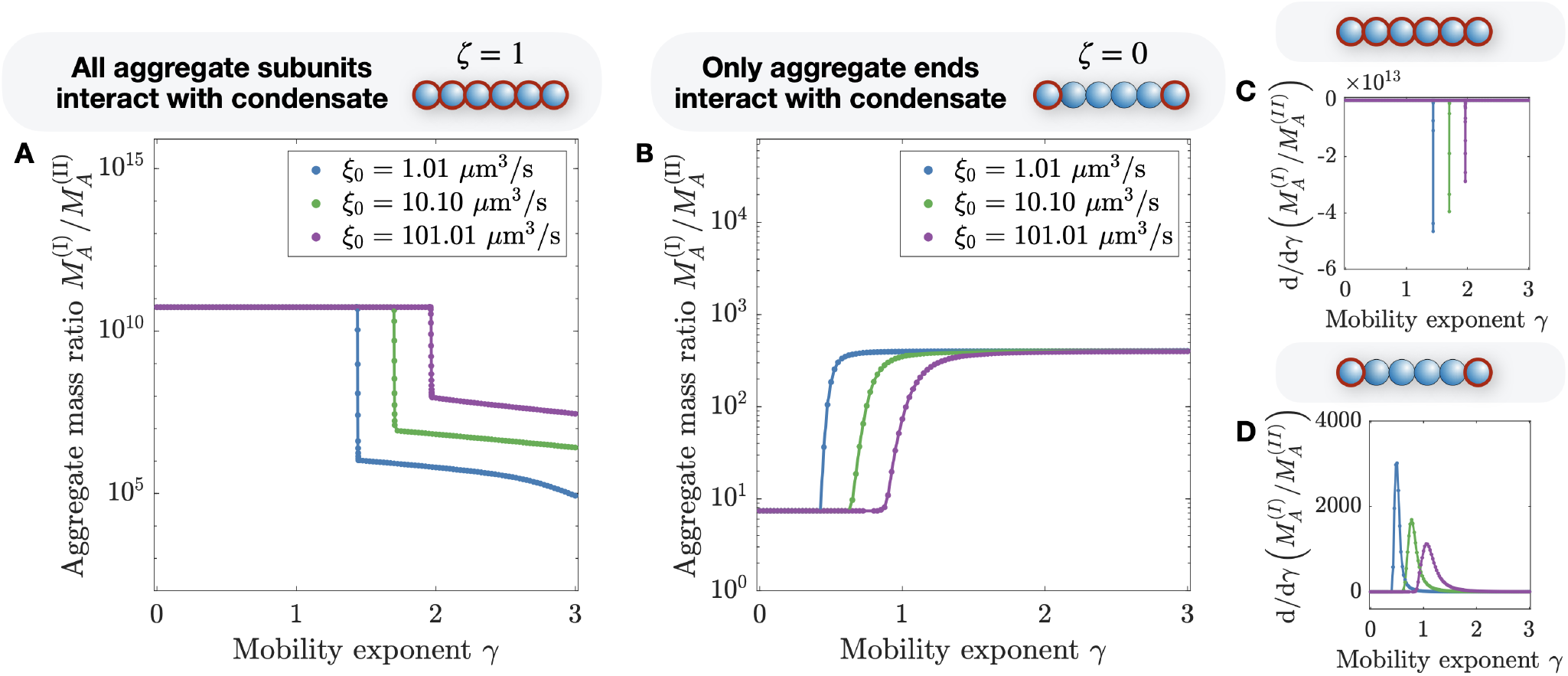
Switch-like transition between different rheology-dependent behaviours of the aggregate mass ratio. For the binding parameter (A) *ζ* = 1 and (B) *ζ* = 0, we show the aggregate mass ratio as a function of the aggregate mobility exponent *γ* for different values of the mobility prefactor *ξ* _0_ (see Eq. (12)). We also calculate the change of the aggregate mass ratio, 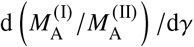 for (C) *ζ* = 1 and (D) *ζ* = 0, The pronounced change of the mass ratio for small variations of the mobility exponent *γ* over multiple orders of magnitudes is reminiscent of a non-equilibrium bifurcation. The total initial monomer concentration is 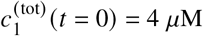 (see Eq. (14)) and the volumes are *V* ^(I)^ = 10.1 *µ*m^3^ and *V* ^(II)^ = 1000 *µ*m^3^. For (A) and we used *P*_*i*_ = exp(2*i*) and for (B) and (D) we used *P*_*i*_ = exp(2).

For *ζ* = 1, the outcome of the aggregation has two limits that we can investigate analytically (see supporting material S1.2): for small values of *γ* and high values of *ξ* _0_, aggregates and monomers are exchanged instantaneously (see supporting material S1.2.2 for the analytical solution). Since the partition coefficient for aggregates, *P*_*i*_, increases with size for *ζ* = 1, aggregates that form outside the condensate are transported into the condensate. For high enough values of *γ*, only monomers are exchanged, while aggregates remain outside or inside of the condensate, depending on where they have formed (see supporting material S1.2.1 for the analytical solution). In Fig. S3, we solve the master equations in these limits numerically and show the excellent agreement with the analytical predictions.

### Properties of switch-like transition when only aggregate ends interact with condensate (*ζ* = 0)

We next investigate the case where only the ends of linear aggregates interact with the condensate (binding parameter *ζ* = 0). In this case, the partition coefficient *P*_*i*_ is independent of aggregate size *i* and a substantial amount of long aggregates is located outside of the condensates since *P*_*i*_ is considerably smaller than in the previous case with *ζ* = 1. Monomers and short linear aggregates form larger aggregates within phase I, leading to an influx from phase II to I of monomers. As a result, aggregates dominantly form within the condensate. The resulting difference in chemical potentials leads to an outflux of aggregates from phase I to II. First, we again use the relation in Eq. (16) to predict the location of the transition in the mass ratio between the inside and outside of the condensate (see Fig. 2 *E*). Then, we investigate and find that near the transition, small changes in the mobility exponent *γ* still lead to big changes in the mass ratio, but less dramatically than for *ζ* = 1.

By comparing the aggregation time and the exchange time *τ* _*f*_, we can predict when exchange of aggregates of size *i* happens on the same time scale as the aggregation for the case of exclusive and instantaneous monomer partitioning ^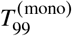^ (see blue lines in Fig. 2 *E*). For aggregate sizes comparable to the front position in the homogeneous case, corresponding to the largest aggregates observed in that case (see Fig. S2), we can recover the location of the observed transition in diagrams (see blue curves in Fig. 2 *E*). Additionally, we observe another transition for values around *>* _0_ = 0.01 − 0.1 *µ*m^3^ /s that can be linked to the exchange of oligomers, *i* = 2.

We next investigate the dependence of the aggregate mass ratio on the mobility exponent *γ*. Analogously to the case with*ζ* = 1, we find two distinct regions in Fig. 2E for *ξ* _0_ *>* 1 *µ*m^3^ /s. By investigating this transition (see Fig. 3 *B*), we find that the transitions happens over a larger range of values of *γ* than for *ζ* = 1 and corresponds to a change of mass ratio of 2 orders of magnitude. Interestingly, the direction of the transition is opposite to the one observed for *ζ* = 1, i.e., increasing the mobility exponent *γ* leads to an increase of the mass ratio. By altering how the condensate is interacting with the linear aggregates, a cell is capable of dramatically altering the outcome of the aggregate distribution in the presence of liquid condensates. For *ξ* _0_ *<* 1 *µ*m^3^ /s, we discover another pronounced transition that is resulting from oligomer exchange (with *i* = 2).

Even though the transition for *ξ* _0_ *>* 1 *µ*m^3^/ s is more smooth, we can still consider two limits between which the transition takes place: For large enough values of *γ* and small enough values of *ξ* _0_, we expect to observe exclusively monomer exchange (see supporting material S1.2.1 for the analytical solution). Importantly, this means that the mass ratio, a quantity that can be accessed experimentally by fluorescently tagging the aggregates and monomers, can exceed the aggregate partition coefficient, that was here chosen to be *P*_*i*_ = exp 2, but reaches values up to 10^4^ for the numerical solutions of the master equations. It is important to note that this results from the rheological features of the condensate. For small values of *γ* and high enough values of *ξ* _0_, all aggregates are rapidly exchanged between both phases and the mass ratio is identical to the partition coefficient (see supporting material S1.2.2 for the analytical solution). But since the partition coefficient is now not dependent on the aggregate size, aggregates generated in phase I will be transported to phase II to fulfill the partition coefficient *P*_*i*_ and the aggregate mass ratio will be smaller and reach *P*_*i*_. This is also consistent with the aggregation time that increases for small *γ* (see Fig. 2 *D*) since the concentration of aggregation within I is decreased, reducing the effects of secondary nucleation and linear aggregate elongation (see Fig. 1 *A*). This has also an effect on the total aggregate number concentration (see Fig. 2 *F*), resulting in a lower aggregate concentration for small values of *γ*. We numerically solve the master equations in both limits and find excellent agreement with the analytical predictions (see Fig. S3 and supporting material S1.2).

### Linear aggregation kinetics in a condensate with complex rheological properties

In the previous section, we considered the case where the aggregate mobility *ξ* _*i*_ is governed by a single power law with the exponent *γ*. If linear aggregates are embedded in a network of phase-separating molecules *A* and *B*, we need to account for diffusion of linear aggregates in a complex liquid environment. For example, condensates composed of reconstituted proteins in-vitro were shown to exhibit a viscoelastic rheology consistent with a Maxwell-like model (35, 50). Moreover, it has been reported that phase-separated biopolymer melts can have pore sizes in the order of 10 nm (51, 52).

A simple model for such complex rheological properties takes into account that the diffusion coefficient follows mixed scalings as a function of aggregate size with multiple crossovers. Here, we study two scaling regimes with one crossover which may present a simple model for a condensate where the phase separating components form pores of characteristic size. In this case, linear aggregates in a condensate follow a Rouse dynamics (mobility exponent *γ* = 1) when they are shorter than the pore size. When aggregates grow in length, they become entangled, for example with the material that forms the condensate. Then, aggregates diffuse similar to polymers in the reptation regime (*γ* = 2) (46, 47, 53). For this simple model with two scaling regimes, the mobility can be written as

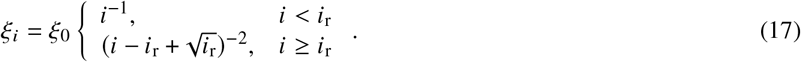

Here, *i*_r_ is the characteristic aggregate size where the transition from Rouse to reptation dynamics takes place, as shown in Fig. 4 *A* and representing a measure of the pore size of the condensate. For *i*_r_ = 1, the mobility is identical to the reptation regime (*γ* = 2), in the limit *i*_r_→∞ the mobility corresponds to the Rouse regime (*γ* = 1).

**Figure 4:**
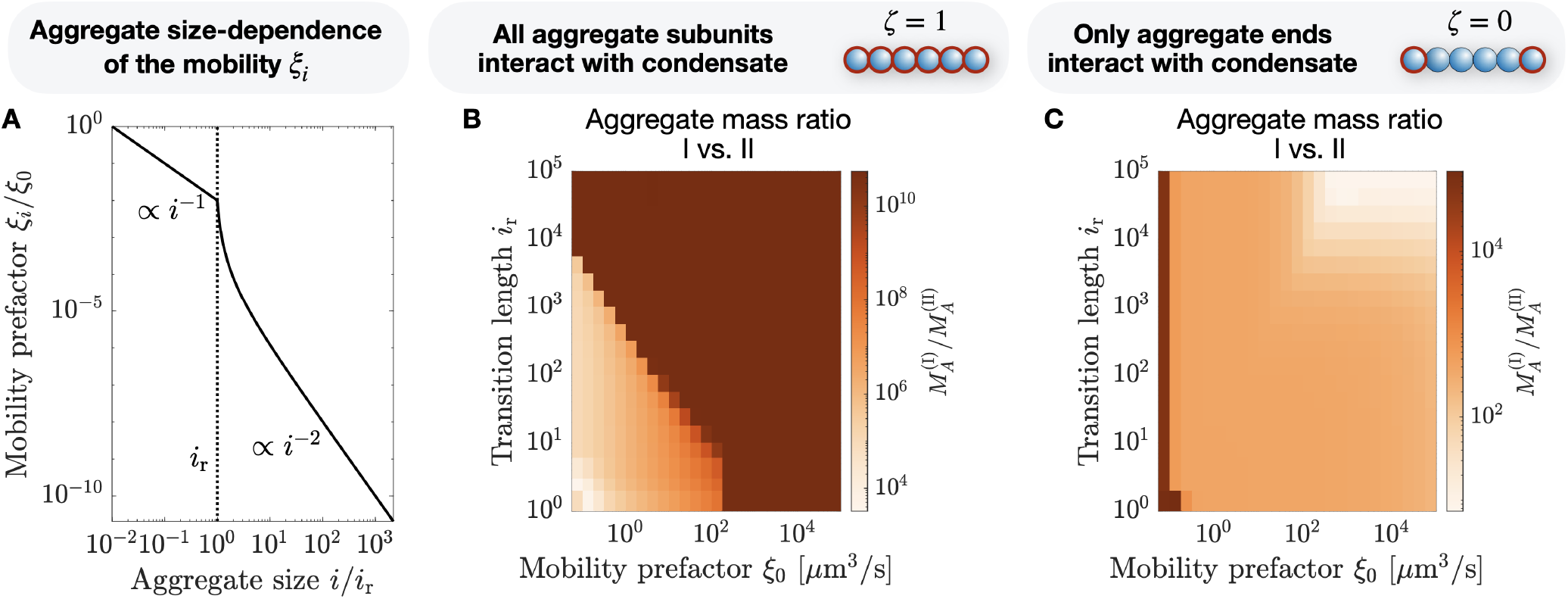
Linear aggregation dynamics for a more realistic dependence of the aggregate diffusion coefficient on aggregate size. (A) Plot of Eq. (17) with the transition length *i*_r_ measured in terms of monomer sizes and mobility prefactor *ξ* _0_. (B) Aggregate mass ratio for *ζ* = 1 with *P*_*i*_ = exp 2*i*. For *i*_r_ = 1 the dynamics corresponds to the reptation regime (*γ* = 2)), for *i*_r_ = ℞ it corresponds to the Rouse regime (*γ* = 1)). We observe a transition between the Rouse and reptation dynamics around *i*_r_ = 10 − 10^4^. (C) Aggregate mass ratio for *ζ* = 0 with *P*_*i*_ = exp 2. We observe a transition between the Rouse and reptation dynamics around *i*_r_ = 1 − 10^3^. The total initial monomer concentration is 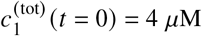 (see Eq. (14)) and the volumes are *V* ^(I)^ = 10.1 *µ*m^3^ and *V* ^(II)^ = 1000 *µ*m^3^.

In Fig. 4 *B* we show the aggregate mass ratio for the case when aggregates bind over their whole length to the phase-separating material A and B, corresponding to the binding parameter *ζ* = 1. Again, we observe a switch-like behaviour, analogously to the behaviour observed in 2 *B*, but now depending on the aggregate length *i*_r_ at which the transition from rouse to reptation dynamics happens instead of the mobility exponent *γ*. While for small enough values of *i*_r_ (for the chosen parameters *i*_r_ *<* 10), the behaviour still corresponds to the reptation regime behaviour. For *i*_r_ *>* 10^4^, the behaviour is again independent of *i*_r_ and corresponds to the Rouse-regime dynamics. This is in accordance with the maximal aggregate size in the system. In between, for 1 *< i*_r_ *<* 10^3^, we see a transition from the reptation regime like behaviour towards the Rouse like behaviour.

If aggregates only bind with their ends to A and B, *ζ* = 0, we see that for the chosen parameters in Fig. 4 *C*, the behaviour is more complex: While again it is independent of the characteristic aggregate size for *i*_r_ *>* 10^4^, for smaller values it undergoes a smooth transition from Rouse (*γ* = 2) to reptation dynamics (*γ* = 1).

We find that a switch-like transition can also occur for condensates of complex rheology by changes of of the transition length *i*_r_. By changing *i*_r_, a cell can control at which mobility the transition occurs. In other words, the characteristic pore size of the condensate material, which should be related to the transition length *i*_r_, affects the aggregation kinetics. Specifically, bigger pores lead to Rouse dynamics with a higher aggregate mass ratio when all aggregate subunits interact with the condensate, *ζ* = 1. If *ζ* = 0, so that aggregates only interact with their ends with the condensate, bigger pores can lead to a smaller mass ratio, as long as the mobility prefactor is large enough.

To summarise, here we provide a numerical study that is as close as possible to experimentally tractable parameters, allowing for a direct comparison to and prediction of experimental studies. This provides us with another way how a cell can control linear aggregation.

## CONCLUSION

We presented a theoretical framework to investigate how the kinetics of irreversible protein aggregation of dilute linear aggregates is affected by the rheological properties of phase-separated condensates. We found that condensate rheology can have a considerable impact on the overall aggregate distribution and aggregation kinetics. In particular, we discovered different regimes of monomer and aggregate transport between the inside and outside of the condensates (see Fig. 5 *A,B*): Either only monomers are exchanged between the two phases on a time scale faster than the aggregation time, or both, monomers and long aggregates. Additionally, we discovered that the way how linear aggregates interact with the phase-separating proteins can have a substantial impact on the distribution of aggregates and how fast monomers assemble (see Fig. 5 *A-C*). A key finding of our work is that slightly different rheological properties, reflected either in different mobility prefactor *ξ* _0_ or mobility exponents *γ* (see Eq. (12)), can lead to pronounced changes of the aggregate distribution. These pronounced changes occur in a very narrow window of controlling parameters indicating switch-like behavior. Due to irreversible kinetics, this transition is not a thermodynamic phase transition rather the transition is reminiscent of a non-equilibrium bifurcation.

**Figure 5:**
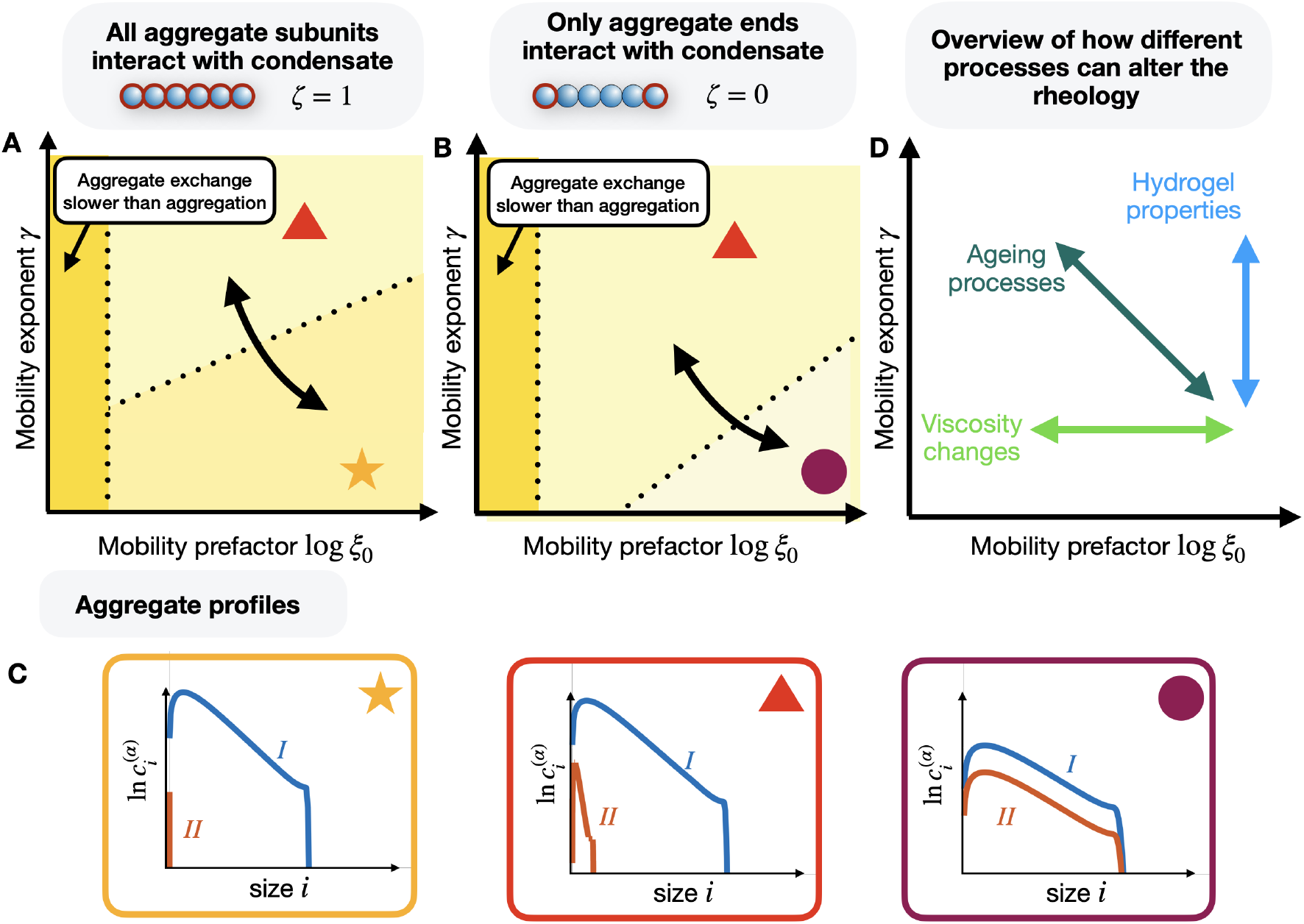
Overview of how condensate rheology can affect linear aggregation. We consider two cases: (A) either all aggregate subunits bind to the condensate, *ζ* = 1, or (B) only aggregate ends bind to the condensate, *ζ* = 0. In both cases, we discover different regimes that affect the aggregate size distribution, depending on the mobility exponent *γ* and prefactor *ξ* _0_. For very small *ξ* _0_, in both cases the exchange of monomers and aggregates is too slow and the aggregation in both phases is independent of each other. For higher values of the mobility prefactor *ξ* _0_ and high values of the mobility exponent *ξ*, in both cases we find that only monomers are exchanged between the compartments. We sketch the resulting aggregate distributions inside (phase I) and outside of the compartment (II) in (C) in the orange box and their location in (A) and (B) with the red triangle. For lower values of the mobility exponent *γ*, aggregates and monomers are exchanged between the inside and outside of the compartment. For *ζ* = 1, this leads to an accumulation of most aggregates inside the condensate. This limit is shown via the orange star in (A) and (C). For *ζ* = 0, the partition coefficient is constant and while aggregates are exchanged, their ratio is fixed. In this case, the accumulation of aggregates inside the condensate is weaker and we can also find a non negligible amount of aggregates outside the condensate. Additionally, the aggregate concentration follows 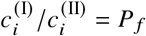 with the fixed aggregate partitioning factor *P* _*f*_. This limit is shown in (B) and (C) with the purple circle. In (A) and (B) we highlight with the black arrows in what direction a cell needs to change its rheology to transition from one limit to another one. In (D) we sketch how different physical processes can alter the rheology of a cellular condensate.

Our theoretical findings of a switch-like change of the aggregation kinetics due to changes in rheology can be experimentally scrutinized by in-vitro studies of droplets undergoing gelation or upon adding agents that affect viscosity. In living systems, various processes can cause changes in rheological properties (see Fig. 5 *D*). One example for such rheological changes are variations of the cytoplasmic concentration via an osmotic shift, which enables cells to alter the aggregation and disassembly of microtubule (54). Recently, it has been discovered that varying the pH level of a cells cytoplasm can mediate a transition between a fluid-like and a solid-like state, referred to as dormant state (34, 55). Thus, a cell possesses a large toolkit that it could use to alter its rheology and as a result, the kinetics of physiological or aberrant aggregates. In the future, unravelling the link between intra-cellular changes in rheology and aggregation kinetics, may pave the way to understand how ageing or the transition to dormancy are linked to protein aggregation-related diseases.

Our theoretical model relies on a set of assumptions. For example, we assumed that linear aggregates are well-mixed inside and outside of the condensates at any time and the flux between the condensate inside and outside is proportional to the chemical potential difference. However, while short aggregates homogenize quickly in each phase due to diffusion and very long aggregates are essentially not exchanged at all, aggregates of intermediate size are expected to transiently accumulate at the phase boundary contradicting a well-mixed assumption. In other words, the concentration of such aggregates may follow a non-linear spatial profile around the interface that presumably decreases towards the center of the condensate. The non-linear profile within the condensate will homogenize on a time-scale that depends on the size of the condensate and aggregates diffusion coefficient. Future steps concern the derivation of the theory governing the time-evolution of size distributions that vary in space. Such a theory enables to investigate the role of heterogeneous size distribution arising from difference in the exchange flux between longer and shorter aggregates. The theory would also enable us to investigate the effect of spatially varying rheological properties. Importantly, we expect that in most cases such effects are only important close to the switch-like transition reported in this work. Another effect that we are ignoring in this study is that in principle, aggregate size could exceed condensate size, potentially leading to kinks and ring-like arrangements of the linear aggregates (56) or to linear filaments emanating from the condensate (57). Capturing such effects also requires a spatial theory for the aggregation kinetics. Additionally, aggregates in experimental systems are not necessarily dilute. Non-dilute aggregation can affect phase separation of condensates. In this case, it is necessary to account for the feedback between phase separation and protein aggregation. Finally, we ignored aggregate fragmentation in this study, since it typically makes the solution of the governing equations significantly more complex. In the future, it will be interesting to study the role of fragmentation, using recently developed mathematical tools (58, 59).

## AUTHOR CONTRIBUTIONS

WP, TM and CW designed the research. WP carried out all numerical solutions. WP and TM carried out analytical calculations. WP analyzed the data. WP, TM and CW wrote the article.

## ACKNOWLEDGMENTS

We thank Giacomo Bartolucci, Sudarshana Laha, Jonathan Bauermann, Guillaume Salbreux and the Salbreux Lab for helpful discussions. WP kindly acknowledges financial support from the Herchel Smith Fund (Herchel Smith Postdoctoral Fellowship). CW acknowledges the financial support by the European Research Council (ERC) under the European Union’s Horizon 2020 research and innovation programme (Grant agreement No. 949021, “FuelledLife”).

## A DETAILS OF NUMERICAL SOLUTION OF THE MASTER EQUATIONS AND ANALYSIS OF CONCENTRATION PROFILES

**Table 1:**
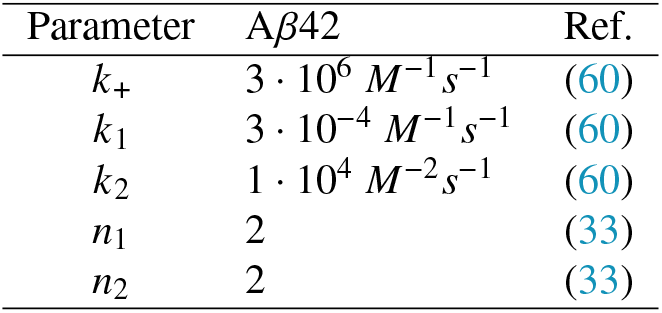
Physicochemcial parameters of A, *β* 42 fibril aggregation estimated by *in-vitro* experiments.

| Parameter | Aβ42 | Ref. |
| --- | --- | --- |
| $k_+$ | $3 \cdot 10^6 \text{ M}^{-1} \text{ s}^{-1}$ | (60) |
| $k_1$ | $3 \cdot 10^{-4} \text{ M}^{-1} \text{ s}^{-1}$ | (60) |
| $k_2$ | $1 \cdot 10^4 \text{ M}^{-2} \text{ s}^{-1}$ | (60) |
| $n_1$ | 2 | (33) |
| $n_2$ | 2 | (33) |

The numerical solution of the master equations (see Eq. (1a) and Eq. (1b)) were performed on the local computing cluster of the MPI-PKS, consisting of X86-64 GNU/Linux systems. The code was written in C++. We used the GCC compiler (version 7.5). In order to numerically solve the master equations (see Eq. (1a) and Eq. (1b)) we used an Euler algorithm to discretize time using a time step *Δ t* = 2 ms.

The data was then analysed with a custom scripts written in Matlab R2020b. The moving front in Fig. S3 *J-L* is identified by first normalising the concentration profiles

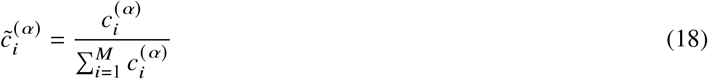

and then only pick the region where *i >* 300 and 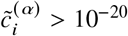 We then compute the logarithm of the concentrations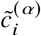 and compute the 100-mean average with the Matlab function movmean(). We find peaks of this curve, corresponding to region with a high concentration gradient, with the function findpeaks() and choose the one that corresponds to the smallest value of *i*.

## B VOLUME FRACTION OF AMYLOID MONOMERS AND FIBRILS

The physiological concentration of amyloid-*β* monomers is around *c*_1_ = 90 − 150 pM (48, 49). The average hydrodynamic radius of a amyloid-*β* is around _1_ = 1 − 3 nm (61–63). The corresponding monomer volume is given by 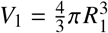 The volume fraction then results from

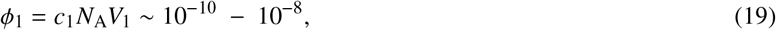

with the Avogadro constant *N*_A_ = 6.022 10^23^ mol^−1^. Thus, the monomers are dilute.

The persistence length of amyloid-*β* fibrils of lengths between 0.2 − 5 *µ*m is around *l*_*p*_ ≈ 4 *µ*m (64). For condensates that are smaller than the persistence length, we approximate aggregates as stiff rods. In that case, the volume of a linear aggregate consisting of *i* monomers is given by

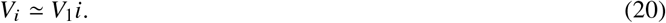

When we consider that monomers of an initial concentration *c*_m_ are all exclusively assembled into aggregates of size *i*, the aggregates have a number concentration of

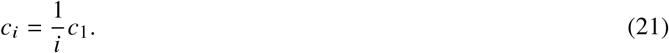

The volume fraction of aggregates consisting each of *i* monomers is then given by (using Eq. (20), Eq. (21) and Eq. (19))

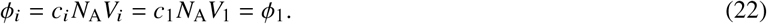

This implies that if monomers are dilute, aggregates will be dilute too. Please note that for condensates that have a diameter smaller than the persistence length, *l*_*p*_ ≈4 *µ*m, the ends of aggregates are not necessarily located within the condensate. In this study, we are ignoring such effects and assume that the aggregates persistence length is smaller than the condensate diameter. If aggregate length is smaller than condensate diameter, we can describe aggregates as spherical aggregates with a volume given by 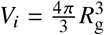 Here, we introduce the radius of gyration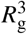. For a good solvent, the radius of gyration follows 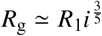 for a bad solvent it follows _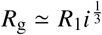_ (46). Thus, the upper limit of the volume of a linear aggregate consisting of *i* monomers (for a good solvent) is given by

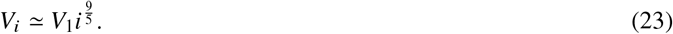

If all monomers with volume fraction *ϕ*_1_ assemble exclusively into aggregates of length *i*, the volume fraction of the aggregates is then given by (using Eq. (23), Eq. (21) and Eq. (19))

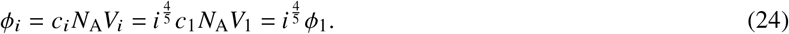

Any linear aggregates assembled by dilute monomers are dilute as long as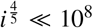. This corresponds to amyloid aggregates consisting of 10^10^ subunits, which exceeds by several of orders the largest aggregates considered in our studies.

## C DERIVATION OF FREE ENERGY

Here, we consider an incompressible mixture of two phase-separating components *A* and *B* with monomers and linear aggregates of different sizes *i*. We assume that the monomers are highly diluted in respect to components of *A* and *B*. The total number 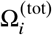 of available states of *N*_*i*_ indistinguishable aggregates of size *i* is given by

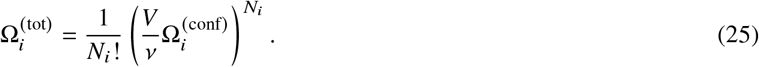

Here, the term *V v* corresponds to the total number of available locations for the aggregate centers of with *V* the system volume and *v* the volume of a monomer. The second term 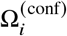 corresponds to the internal configurations of the aggregates, which is for example affected by whether the aggregate is located in a good or poor solvent. Since the linear aggregates are dilute, we assume that each of them can be independently distributed in the volume *V* and there are no excluded volume effects, so that their internal configurations are not affected by the presence of other aggregates. Additionally, we assume that the presence of components *A* and *B* has no effect on 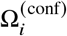. The mixing entropy for aggregates of size *i* then follows from

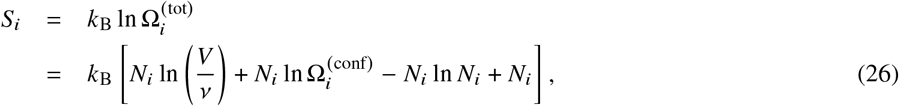

where we have used Sterling’s approximation ln (*N*!) ≈ *N* ln *N* − *N*. The total mixing entropy, also taking into account the components *A* and *B* and aggregates of size *i* ∈ {1, …, *M* }, is then given by

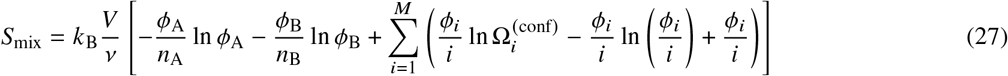

with the linear aggregate volume fractions *ϕ*_*i*_ = *iεN*_*i/*_ *V* of aggregates of size *i*, the component volume fractions *ϕ*_A_ and *ϕ*_B_, with *n*_A_ and *n*_B_, the non-dimensional sizes of components *A* and *B* in multiples of *ν* and the maximal aggregate size *M*. Since we assume that *A* and *B* are not dilute, we can us to use the same mixing entropy contributions as in the Flory-Huggins theory.

Next to the mixing entropy, the enthalpy due to interactions of the aggregates with the components *A* and *B* also contributes to the free energy. For *N*_*i*_ aggregates of size *i*, the enthalpy is given by

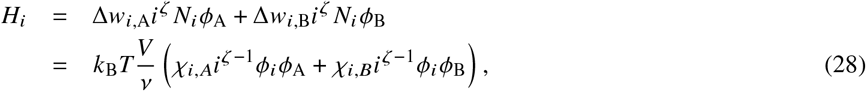

where Δ*w*_*i*,A_ and Δ*w*_*i*,B_ are the change of energy per monomer of the linear aggregate in contact with components *A* and *B* and defining the interaction parameters *χ*_*i,A*_ = Δ*w*_*i*,A_ /*k*_B_ and *χ*_*i,B*_ = Δ*w*_*i*,B_ *k*_B_. Here, the binding parameter *ζ* represents the limit of how the phase separating components *A* and *B* interact with the aggregate: for *ζ* = 1 over its full length and for *ζ* = 0 only with the two linear aggregate ends (see Fig. 1 *C*). For *ζ* = 1, the enthalpy term is identical to the one found for the Flory-Huggins theory since we can still make the mean field approximation for the components *A* and *B*.

The total enthalpy of the system is then given by

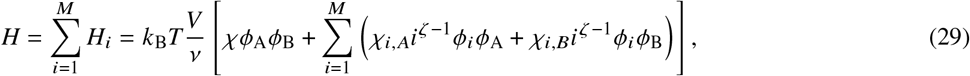

ignoring interactions of the aggregates with each other since we assume that they are far apart from each other and do not intersect. The complete free energy is then given by

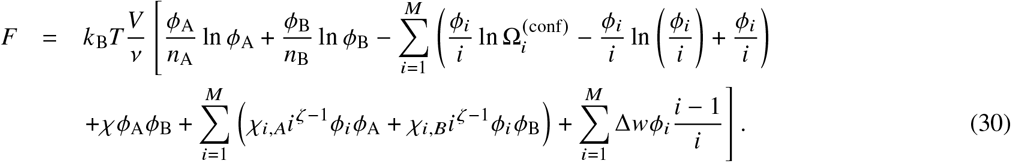

Here, we also introduced the bond energy f..*w* between each subunit pair bound to each other within an aggregate. For an aggregate of length *i*, there are *i* − 1 subunits pairs. Since the linear aggregates are highly dilute, we can assume that the volume fractions of the components can be approximated by *ϕ*_A_ = and *ϕ*_B_ = 1 and 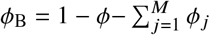 so that the total free energy of the system reads

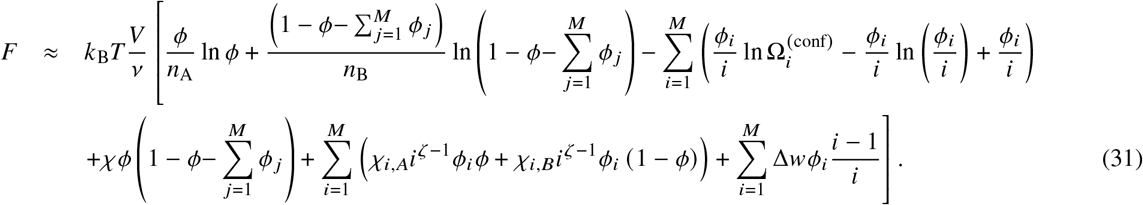

## D AGGREGATION KINETICS IN COEXISTING PHASES

For simplicity, we will now consider the special case of monomer pickup and investigate how phase equilibrium affects the aggregation dynamics. The calculations are based on Ref. (38). The chemical potential given in Eq. (7) can be written as

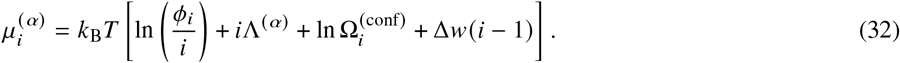

We introduce the phase-dependent term (*α* = I, II)

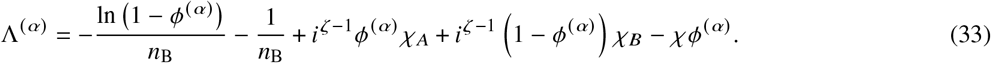

For monomer pickup (see Fig. 1A), we consider an irreversible process with the rate

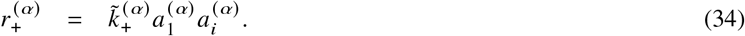

Here, we introduce the chemical activities 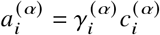 with the activity coefficients 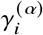 The rate can be written as

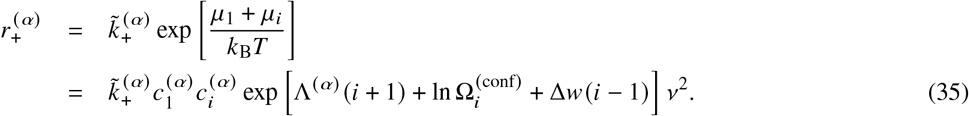

where we use a relationship between the activities and the chemical potentials of the form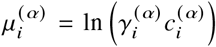, which corresponds to the case of zero reference chemical potentials. The activity coefficients are

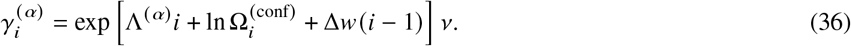

As shown in Ref. (38), the activity coefficients satisfy

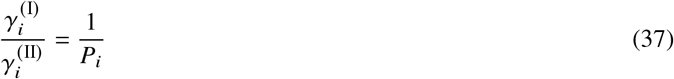

at phase equilibrium.

The interaction parameters *χ, χ*_*A*_ and *χ*_*B*_ typically have amplitudes of a few *k*_B_ (33, 34), while the magnitude of the bond energyΔ*w* is much bigger since we consider irreversible aggregation. In this case, we can approximately write

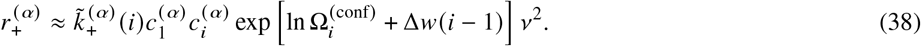

When the coexisting phases are liquids, only the rate constants 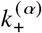 and the concentrations depends on the respective phases. In the homogeneous case, the reaction rate reads

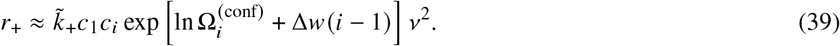

According to previous experimental studies of homogeneous systems (14, 15, 17, 37), there is no significant dependence of the reaction rate on aggregate length *i*, i.e.,

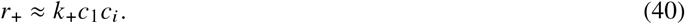

To be consistent with these experimental measurements, we choose the reaction rate constant 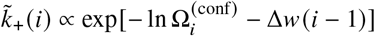. Analogously, we obtain for the case with two phases

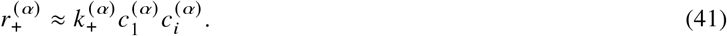

Analog steps can be performed for the nucleation processes, finally leading to Eqs. (1).

## E RELATIONSHIP BETWEEN AGGREGATE MOBILITY AND AGGREGATE DIFFUSION COEFFICIENT

To derive the relation between the mobility coefficient *ξ*_*i*_ of aggregates of size *i* (see Eq. (12)) and the corresponding diffusion coefficient *D*_*i*_, we consider weak concentration profiles of the concentrations of aggregates. In this case, the mobility in each phase is approximately constant. Moreover, we focus on the limiting case where the reaction kinetics is slower than the exchange kinetics between the phases suggesting a quasi static approximation for the diffusive exchange between the phases. For such quasi static conditions, the concentration profile is determined by the Laplace equation,

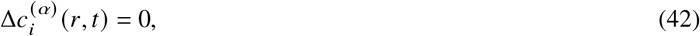

where time characterizes the slow transient due to the aggregation kinetics. For simplicity, we solve the Laplace equation for small spherical condensed phases with radius in a finite systems of size *L*, where *R* « *L*. For slow reaction steps and small enough condensates, concentration gradients within the condensate 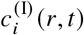 can be neglected compare to the surrounding phase, i.e.,

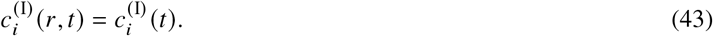

For the surrounding phase II, we find approximately (approximation refers to neglecting terms of the order /*L*)

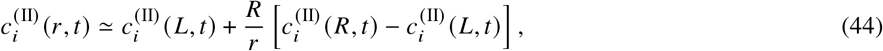

where 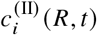 denotes the concentration directly at the interface of phase II. Applying partition equilibrium, we obtain

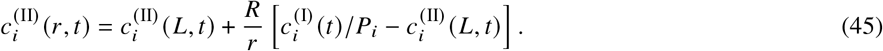

Next, we the radial flux 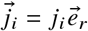 is given by

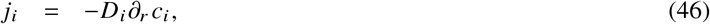

where *D*_*i*_ denotes the diffusion coefficient of aggregate of size *i*. At the interface *r*=*R*, the radial flux reads

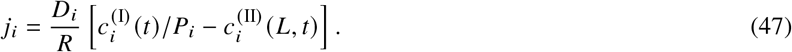

The total exchange rate _*i*_ [units: 1/s] (Eq. (11)) is related to the radial flux by _*i*_ = 4 π^2^ *j*_*i*_ leading to

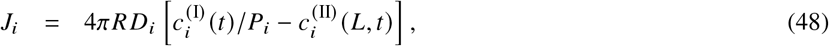

Comparing to linearization of Eq. (11) (see also Ref. (33)), we find the following relationship between the diffusion coefficient and the mobility:

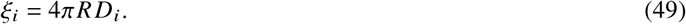

This relationships allows us to estimate the range of diffusion coefficients that correspond to the range of mobilities considered in our work. Please note that this estimate depends on the condensate radius *R*. We consider typical values of condensate radii, i.e., *R* = 0.1, 1, 10 *µm*. For amyloid monomers, reported diffusion coefficient is around *D*_0_ ≈ 100 *µm*^2^ s (63). For the condensate radii given above, the corresponding mobilites are *ξ*_0_ = {10^2^, 10^3^, 10^4}^ *µm* ^3^ *s*. In our simulations, we consider mobility values in this range as well as a bit higher and lower for the sake of generality. Thus, we probe experimentally relevant parameter values for the mobilties.

## F CHARACTERISTIC TIME OF LINEAR AGGREGATE EXCHANGE BETWEEN PHASES

In equation (11), we give the flux _*i*_ of linear aggregates of size *i* between two coexisting phases I and II. Assuming that 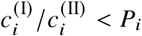, the dynamic equations of the aggregate concentration are written as

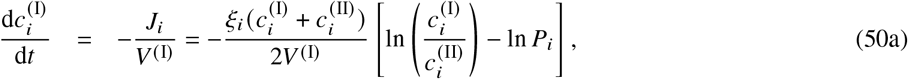

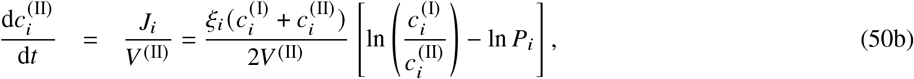

with the partition coefficient *P*_*i*_. In our system, we usually find 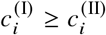. This implies that ln 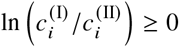. If 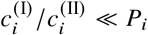 we can then simplify the equations and get

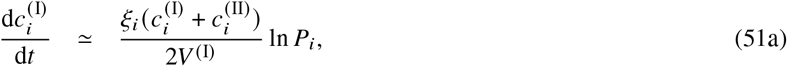

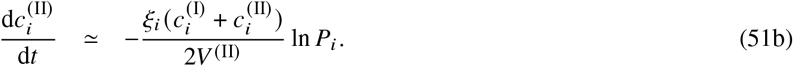

The solution of the equations has a single characteristic time-scale

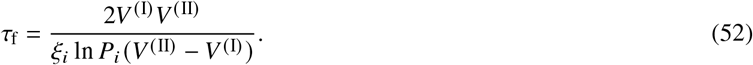

For *V* ^(II)^ » *V* ^(I)^, we find

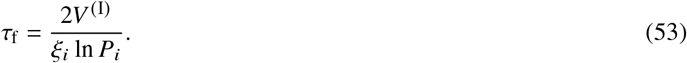

## SUPPORTING MATERIAL

See attached file.

