## Supporting Material for "Aggregation controlled by condensate rheology"

June 14, 2022

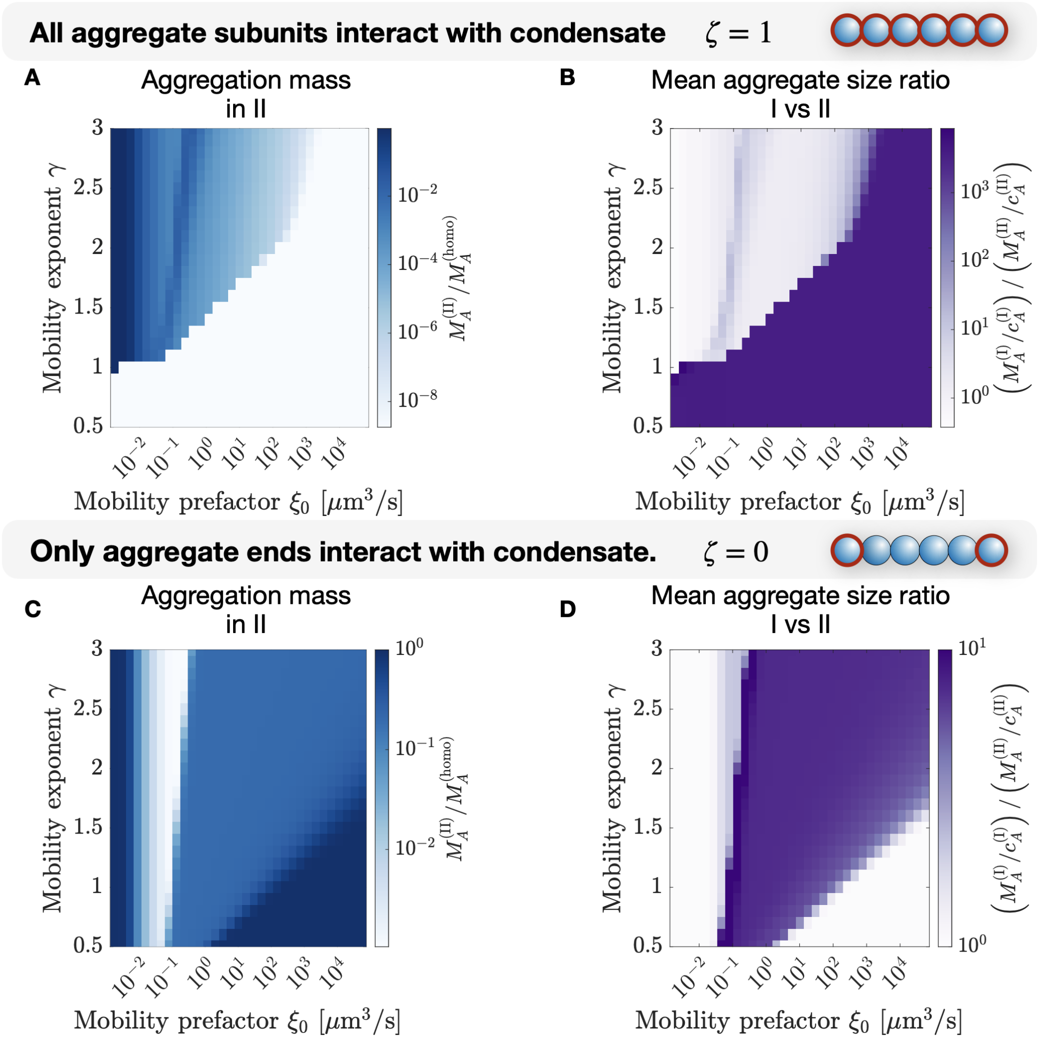


Figure S1: **Diagrams of how droplet rheology affects aggregate mass outside of the droplet and the size ratio of linear aggregates** For the two cases when all aggregate subunits interact with the condensate, ($\zeta = 1$, the partitioning coefficient is given by $P_i = \exp 2i$) or when only aggregate ends interact with the condensate ($\zeta = 0$, the partitioning coefficient $P_i = \exp 2$), we solve the master equation (see Eq. (1a) and Eq. (1b)) and here show aggregate mass concentration in compartment II after the time it took to assemble 99% monomers for (A) $\zeta = 1$ and (C) $\zeta = 0$. The mass concentration is normalised by mass concentration $M_{\mathrm{A}}^{(\mathrm{homo})}$ of the homogeneous case with only a single compartment with the same initial total monomer concentration. In (B,D), we provide the aggregate size ratio, where the size in compartment $\alpha = I, II$ results from $M_{\mathrm{A}}^{(\alpha)}/c_{\mathrm{A}}^{(\alpha)}$. For the solution of the master equation, we chose a total initial monomer concentration of $c_1^{(\mathrm{tot})}(t=0) = 4\ \mu$M (see Eq. 13) and the volumes are $V^{(\mathrm{I})} = 10.1\ \mu\mathrm{m}^3$ and $V^{(\mathrm{II})} = 1000\ \mu\mathrm{m}^3$.

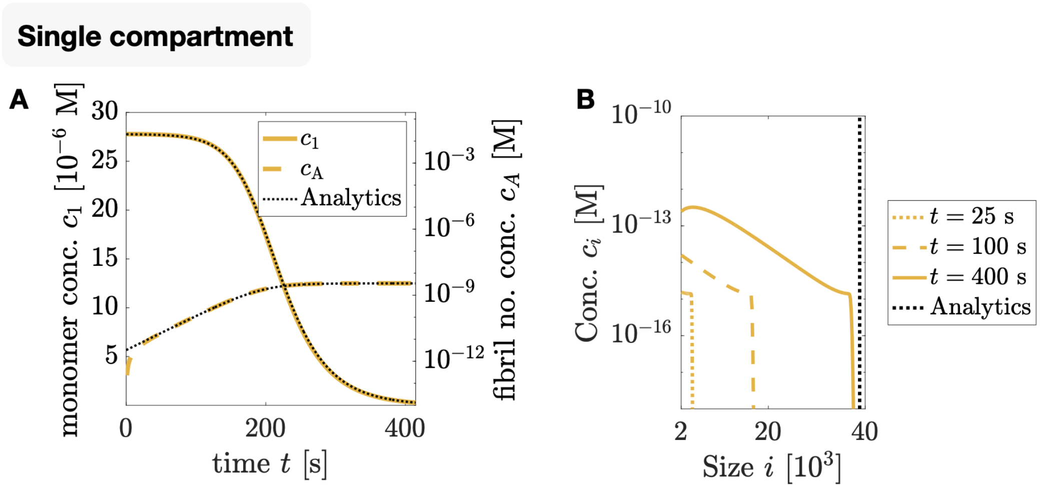


Figure S2: **Linear aggregate concentrations for aggregation in a homogeneous system** (A) Monomer concentration $c_1$ (solid line) and aggregate number concentration $c_A$ (dashed line, see Eq. (2)) from the numerical solution of the master equation (Eq. (1a) and Eq. (1b)). Here, initially only monomers exist with a concentration $c_1(t = 0) = 27.78 \times 10^{-6}$ M to mimick a higher concentration within a droplet. The numerical solution is in excellent agreement with analytical predictions , see Eq. (S8) and Eq. (S9) (black dotted line). While the monomer concentration is decaying with time due to aggregation, the number of aggregates is increasing, until it saturates since almost all monomers are gone and primary nucleation no longer creates new fibrils. (B) Concentration profiles as a function of time. The profile possesses a moving front where the concentration decreases sharply. The black dotted line is the analytical prediction of the front position at the equilibrium, derived in Eq. 42.

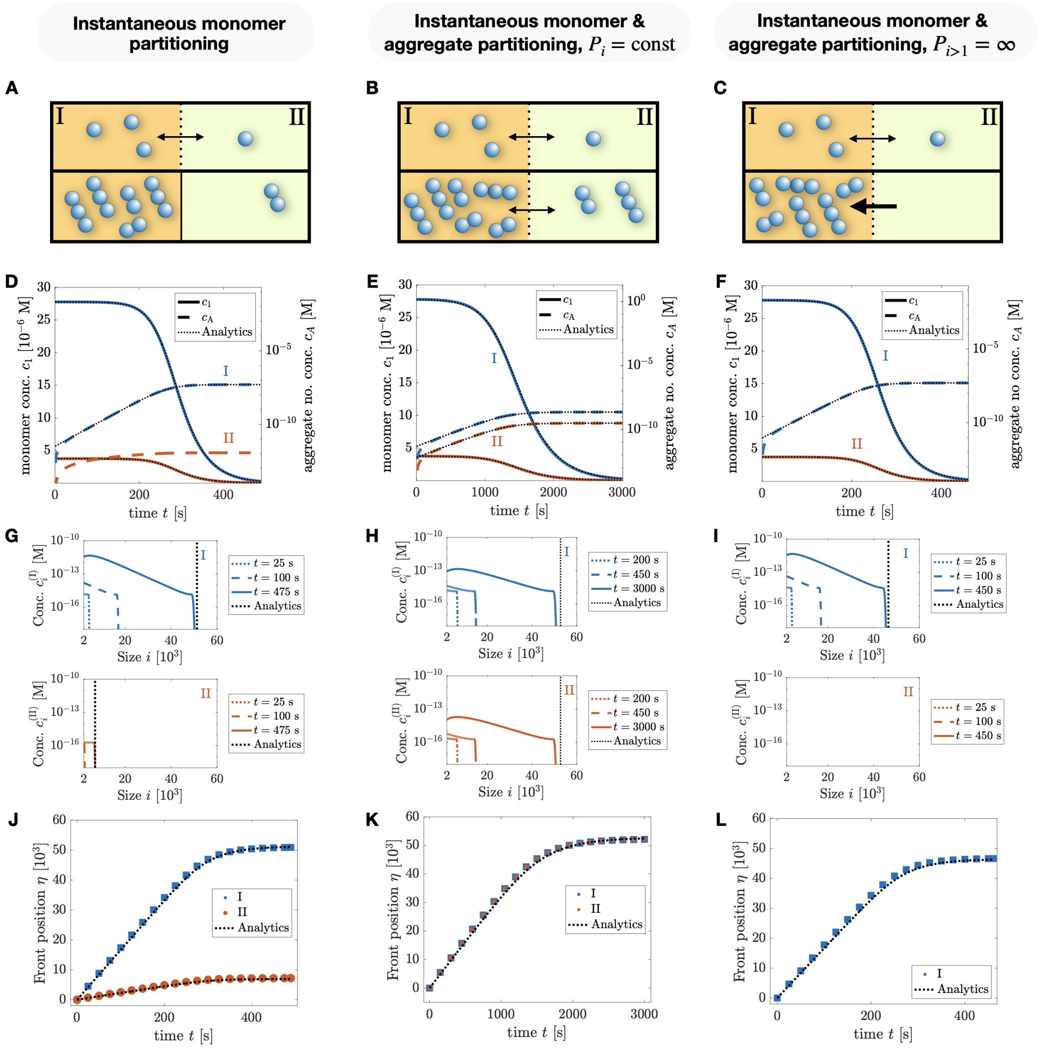


Figure S3: **Comparison of numerical and analytical solution for three different limits of instantaneous monomer and aggregate exchange** The three limits we are considering are: (A) Exclusive monomer exchange, (B) monomer and aggregate exchange with constant partitioning coefficient $P_i = \exp(2)$ and (C) monomer and aggregate exchange with monomer partitioning coefficient $P_1 = \exp(2)$ and aggregate partitioning coefficient $P_{i>1} = \infty$. In (D-F) we show the numerical solutions of the master equation (Eq. (1a) and Eq. (1b)) for the monomer concentration $c_1$ and the aggregate number concentration $c_\mathrm{A}$ for each compartment. The solutions are in excellent agreement with the analytical predictions, provided in Section S1.2.2 and Section S1.2.3.

Figure S3: (G-I)Concentration profiles as a function of time for the three cases in both compartments I (inside the droplet) and II (outside the droplet). The profiles possess a front where the concentrations decay abruptly. This front emerges from the fact that the aggregate assemble for a finite time and larger aggregates did not yet have time to form. The black dotted line corresponds to the position of the front in equilibrium, as predicted analytically. (J-L) Moving front position as a function of time from the numerical solution. How the front position is derived from the concentration profile is explained in Appendix A. he black dotted line corresponds to the analytical predictions. The total initial monomer concentration is $c_1^{(\mathrm{tot})}(t = 0) = 4\ \mu\mathrm{M}$ (see Eq. (14)) and the volumes are $V^{(\mathrm{I})} = 10.1\ \mu\mathrm{m}^3$ and $V^{(\mathrm{II})} = 1000\ \mu\mathrm{m}^3$.

### S1 Analytical solutions for instantaneous exchange of monomers and linear aggregates

Before we derive the analytical solution of the linear aggregation of monomers and aggregates instantaneously exchanged between two phases, we first give the solutions for the single phase case, as previously derived in [1]. Then, we solve the analytical cases of exclusive and instantaneous monomer exchange, on time-scales much faster than the aggregation time-scale [2]. Finally, we solve the case with both, instantaneous monomer and aggregate exchange, and investigate the limit of infinite aggregate partitioning coefficient $P_\mathrm{f}$.

#### S1.1 Single phase

**Constitutive equations** In a homogeneous system, with only a single phase, the aggregation dynamics is described by the equations

$$\begin{aligned}\frac{\mathrm{d}c_1(t)}{\mathrm{d}t} &= -2k_+c_1(t)c_\mathrm{a}(t) \\ &\quad -k_1n_1c_1(t)^{n_1} - k_2n_2c_1(t)^{n_2}M_\mathrm{a}(t) \qquad \text{(S1a)}\\ \frac{\mathrm{d}c_\mathrm{a}(t)}{\mathrm{d}t} &= k_1c_1(t)^{n_1} + k_2c_1(t)^{n_2}M_\mathrm{a}(t) \qquad \text{(S1b)}\end{aligned}$$

for the monomer concentration $c_1(t)$, the aggregate number concentration $c_\mathrm{a}(t)$ and the aggregate mass concentration $M_\mathrm{a}(t) = M_\mathrm{tot} - c_1(t)$. For typical filamentous systems, growth by elongation is much faster than primary and secondary nucleation [1]. In this regime, Eq. (S1a) and (S1b) can be simplified to

$$\begin{aligned}\frac{\mathrm{d}c_1(t)}{\mathrm{d}t} &= -2k_+c_1(t)c_\mathrm{a}(t)\ , \qquad \text{(S2a)}\\ \frac{\mathrm{d}c_\mathrm{a}(t)}{\mathrm{d}t} &= k_1c_1(t)^{n_1} + k_2c_1(t)^{n_2}M_\mathrm{a}(t) \qquad \text{(S2b)}\end{aligned}$$

where we have neglected nucleation terms in the equation for $\mathrm{d}c_1/\mathrm{d}t$. The initial conditions for Eq. (S2a) and (S2b) are $c_1(t=0) = M_\mathrm{tot}$ and $c_\mathrm{a}(t = 0) = 0$.

**Analytical solutions for the monomer and aggregate number concentrations** Equations (S2) has been solved analytically in [1] by using an analogy to classical mechanics, rewriting the system of differential equations as

$$\frac{\mathrm{d}q}{\mathrm{d}t} = \frac{\partial \mathcal{H}}{\partial p}, \tag{S3a}$$

$$\frac{\mathrm{d}p}{\mathrm{d}t} = -\frac{\partial \mathcal{H}}{\partial q}, \tag{S3b}$$

with

$$p(t) = 2k_{+}c_{\mathrm{a}}(t), \tag{S4a}$$

$$q(t) = \ln\left(\frac{M_{\mathrm{tot}}}{c_1(t)}\right), \tag{S4b}$$

$$\mathcal{H}(p,q) = \frac{p^2}{2} + V(q), \tag{S4c}$$

and the 'potential energy'

$$V(q) = \lambda^2 \frac{\exp(-n_1 q)}{n_1} + \kappa^2 \frac{\exp(-n_2 q)\left[(n_2+1) - n_2 \exp(-q)\right]}{n_2(n_2+1)}. \tag{S5}$$

Here, we introduce the parameters

$$\lambda = \sqrt{2k_{+}k_1 M_{\mathrm{tot}}^{n_1}}, \tag{S6a}$$

$$\kappa = \sqrt{2k_{+}k_2 M_{\mathrm{tot}}^{n_2+1}}, \tag{S6b}$$

$$\theta = \sqrt{\frac{2}{n_2\left(n_2+1\right)}}, \tag{S6c}$$

$$\omega = \frac{\lambda^2}{2\kappa^2\theta}, \tag{S6d}$$

assuming that at time $t = 0$, only monomers exist with the concentration $c_1(0) = M_{\mathrm{tot}}$. Using the mass conservation

$$M_{\mathrm{a}}(t) = c_1(0) - c_1(t), \tag{S7}$$

and solving the Euler-Lagrange equation $\frac{\mathrm{d}^2 q}{\mathrm{d}t^2} = -\partial_q V$, we can finally derive an approximate solution for the monomer concentration, given by (see Fig. S2*A*)

$$c_1(t) = M_{\mathrm{tot}}\left[1 + \omega \exp\left(\kappa t\right)\right]^{-\theta}. \tag{S8}$$

The aggregate number concentration $c_\mathrm{a}(t)$ results from Eq. (S2) and Eq. (S8) and is given by (see Fig. S2 *A*)

$$\begin{aligned} c_\mathrm{a}(t) &= -\frac{1}{2k_+ c_1(t)}\frac{\mathrm{d}c_1}{\mathrm{d}t} \\ &= \frac{\theta\kappa}{2k_+}\frac{\omega\exp(\kappa t)}{1+\omega\exp(\kappa t)}, \end{aligned} \tag{S9}$$

which for $t\to\infty$ converges towards

$$\lim_{t\to\infty} c_\mathrm{a}(t) = \frac{\theta\kappa}{2k_+} \propto M_\mathrm{tot}^{\frac{n_2+1}{2}}. \tag{S10}$$

From Eq. (S8) we find that the characteristic rate of aggregation is given by

$$\kappa = \sqrt{2k_+k_2M_\mathrm{tot}^{n_2+1}} \propto M_\mathrm{tot}^{\frac{n_2+1}{2}}. \tag{S11}$$

**Aggregation time** Using Eq. (S8) and assuming that initially there are only monomers, after the time

$$T_\mathrm{p}^{(\mathrm{homo})} = \frac{1}{\kappa}\ln\left(\frac{(1-p)^{-\frac{1}{\theta}}-1}{\omega}\right) \tag{S12}$$

a fraction $p$ of monomers have assembled. For example, if 99% of monomers have been used up, we set $p=0.99$.

**Moving front position** We now introduce the variable

$$\eta(t) = \int_0^t \mathrm{d}t'\; 2k_+c_1(t') \tag{S13}$$

which corresponds to the average number of monomers used up by the growth of one aggregate at time $t$ and hence is a measure of the largest linear aggregate size observed in the system. It is also a measure of the size for which the aggregate concentration profile is sharply declining, called the front (see Fig. S2 *B*). $\eta$ can be calculated using Eq. (S9). The resulting expression is in general complicated, involving hypergeometric functions. We can analytically solve it for $\theta = 0.5$, corresponding to $n_2 \approx 2.37$. Then, the front position $\eta$ is given by

$$\eta(t) = \frac{4k_+ M_{\mathrm{tot}}}{\kappa}\left[\tanh^{-1}\left(\sqrt{1+\omega}\right) - \tanh^{-1}\left(\sqrt{1+\omega\exp(\kappa t)}\right)\right], \tag{S14}$$

$$\eta(\infty) = \frac{4k_+ M_{\mathrm{tot}}}{\kappa}\sinh^{-1}\left(\sqrt{\frac{1}{\omega}}\right). \tag{S15}$$

#### S1.2 Partitioning

We next consider the case where linear aggregates undergo aggregation within the phases I and II, but are also exchanged instantaneously between the phases by a total flux (see Eq. (11)). This corresponds to very high mobilities $\xi_0$ for monomers and $\xi_{\mathrm{f}}$ for aggregates.

**Constitutive equations** The dynamic equations of the monomer and aggregate number and mass concentrations, adapted from [2], are given by

$$\frac{\mathrm{d}c_1^{(\mathrm{I})}(t)}{\mathrm{d}t} = -2k_+ c_1^{(\mathrm{I})}(t)c_\mathrm{a}^{(\mathrm{I})}(t) - \xi_0\left[c_1^{(\mathrm{I})}(t) - Pc_1^{(\mathrm{II})}(t)\right]\frac{1}{V^{(\mathrm{I})}}, \tag{S16a}$$

$$\frac{\mathrm{d}c_1^{(\mathrm{II})}(t)}{\mathrm{d}t} = -2k_+ c_1^{(\mathrm{II})}(t)c_\mathrm{a}^{(\mathrm{II})}(t) + \xi_0\left[c_1^{(\mathrm{I})}(t) - Pc_1^{(\mathrm{II})}(t)\right]\frac{1}{V^{(\mathrm{II})}}, \tag{S16b}$$

$$\frac{\mathrm{d}M_\mathrm{a}^{(\mathrm{I})}(t)}{\mathrm{d}t} = 2k_+ c_1^{(\mathrm{I})}(t)c_\mathrm{a}^{(\mathrm{I})}(t) - \xi_\mathrm{f}\left[M_\mathrm{a}^{(\mathrm{I})}(t) - P_\mathrm{f}M_\mathrm{a}^{(\mathrm{II})}(t)\right]\frac{1}{V^{(\mathrm{I})}}, \tag{S16c}$$

$$\frac{\mathrm{d}M_\mathrm{a}^{(\mathrm{II})}(t)}{\mathrm{d}t} = 2k_+ c_1^{(\mathrm{II})}(t)c_\mathrm{a}^{(\mathrm{II})}(t) + \xi_\mathrm{f}\left[M_\mathrm{a}^{(\mathrm{I})}(t) - P_\mathrm{f}M_\mathrm{a}^{(\mathrm{II})}(t)\right]\frac{1}{V^{(\mathrm{II})}}, \tag{S16d}$$

$$\frac{\mathrm{d}c_\mathrm{a}^{(\mathrm{I})}(t)}{\mathrm{d}t} = k_1 c_1^{(\mathrm{I})}(t)^{n_1} + k_2 c_1^{(\mathrm{I})}(t)^{n_2} M_\mathrm{a}^{(\mathrm{I})}(t) - \xi_\mathrm{f}\left[c_\mathrm{a}^{(\mathrm{I})}(t) - P_\mathrm{f}c_\mathrm{a}^{(\mathrm{II})}(t)\right]\frac{1}{V^{(\mathrm{I})}}, \tag{S16e}$$

$$\frac{\mathrm{d}c_\mathrm{a}^{(\mathrm{II})}(t)}{\mathrm{d}t} = k_1 c_1^{(\mathrm{II})}(t)^{n_1} + k_2 c_1^{(\mathrm{II})}(t)^{n_2} M_\mathrm{a}^{(\mathrm{II})}(t) + \xi_\mathrm{f}\left[c_\mathrm{a}^{(\mathrm{I})}(t) - P_\mathrm{f}c_\mathrm{a}^{(\mathrm{II})}(t)\right]\frac{1}{V^{(\mathrm{II})}} \tag{S16f}$$

where we introduced the aggregate partitioning coefficient $P_\mathrm{f}$, mobility $\xi_\mathrm{f}$, the monomer concentration $c_1^{(\alpha)}$, the aggregate number concentration $c_\mathrm{a}^{(\alpha)}$ and the aggregate mass concentration $M_\mathrm{a}^{(\alpha)}$, for $\alpha = \mathrm{I}, \mathrm{II}$. Here, we linearized the fluxes (see Eq. (11)). The sole task of the fluxes in the constitutive equations is to enforce the ratios of the number and mass concentrations ($c_1^{(\alpha)}$, $c_\mathrm{a}^{(\alpha)}$ and $M_\mathrm{a}^{(\alpha)}$) are equal to the partitioning coefficients $P$ and $P_\mathrm{f}$. Thus, the exact form of the fluxes has no impact on the analytical solutions.

**Initial condition and mass conservation** Here, and in the following, we will assume that initially only monomers exist with

$$c_1^{(\mathrm{I})}(0)V^{(\mathrm{I})} + c_1^{(\mathrm{II})}(0)V^{(\mathrm{II})} = M_\mathrm{tot}\left(V^{(\mathrm{I})} + V^{(\mathrm{II})}\right), \tag{S17}$$

where we introduce the total mass concentration $M_\mathrm{tot}$. Together with the assumption that $c_1^{(\mathrm{I})}(0) = Pc_1^{(\mathrm{II})}(0)$ we get

$$c_1^{(\mathrm{I})}(0) = M_\mathrm{tot}\frac{V^{(\mathrm{I})} + V^{(\mathrm{II})}}{V^{(\mathrm{I})} + \frac{1}{P}V^{(\mathrm{II})}}, \tag{S18a}$$

$$c_1^{(\mathrm{II})}(0) = \frac{1}{P}M_\mathrm{tot}\frac{V^{(\mathrm{I})} + V^{(\mathrm{II})}}{V^{(\mathrm{I})} + \frac{1}{P}V^{(\mathrm{II})}}. \tag{S18b}$$

Mass conservation leads to

$$M_\mathrm{a}^{(\mathrm{I})}(t) = \frac{c_1^{(\mathrm{I})}(0)V^{(\mathrm{I})} + c_1^{(\mathrm{II})}(0)V^{(\mathrm{II})} - c_1^{(\mathrm{I})}(t)V^{(\mathrm{I})} - \left(c_1^{(\mathrm{II})}(t) + M_\mathrm{a}^{(\mathrm{II})}(t)\right)V^{(\mathrm{II})}}{V^{(\mathrm{I})}}, \tag{S19}$$

with the phase volumes $V^{(\mathrm{I})}$ and $V^{(\mathrm{II})}$.

We now provide analytical solutions to the constitutive equations in three limits of practical importance.

##### S1.2.1 No transport between phases I and II

If partitioning between the phases is prohibited, corresponding to $\xi_0 = 0$ and $\xi_\mathrm{f} = 0$, in each phase the aggregation is equivalent to the aggregation dynamics in the homogeneous case (see Section S1.1) with the initial monomer concentrations $c_1^{(\mathrm{I})}(0)$ in phase I and $c_1^{(\mathrm{II})}(0)$ in phase II.

##### S1.2.2 Rapid transport of monomers between two phases

This case has been previously investigated in [2] and is sketched in Fig. S3 *A*.

**Assumptions** Here, we assume that only monomers are transported on such a short time-scale, that at any time the condition

$$c_1^{(\mathrm{I})}(t) = P c_1^{(\mathrm{II})}(t) \tag{S20}$$

is fulfilled. There is no exchange of aggregates between the two phases. Additionally, we choose the initial conditions and $P$ such that within phase II the monomer concentration is continuously smaller than in phase I, so that the aggregate mass and number concentrations in II are negligible, $M_\mathrm{a}^{(\mathrm{II})} \approx 0$ and $c_\mathrm{a}^{(\mathrm{II})} \approx 0$.

**Analytical solutions for the monomer and aggregate number concentrations** The total monomer concentration in the system follows from

$$c_1(t) = \frac{V^{(\mathrm{I})}}{V^{(\mathrm{I})}+V^{(\mathrm{II})}} c_1^{(\mathrm{I})}(t) + \frac{V^{(\mathrm{II})}}{V^{(\mathrm{I})}+V^{(\mathrm{II})}} c_1^{(\mathrm{II})}(t), \tag{S21}$$

$$\frac{\mathrm{d}c_1(t)}{\mathrm{d}t} = \frac{V^{(\mathrm{I})}}{V^{(\mathrm{I})}+V^{(\mathrm{II})}} \frac{\mathrm{d}c_1^{(\mathrm{I})}(t)}{\mathrm{d}t} + \frac{V^{(\mathrm{II})}}{V^{(\mathrm{I})}+V^{(\mathrm{II})}} \frac{\mathrm{d}c_1^{(\mathrm{II})}(t)}{\mathrm{d}t}. \tag{S22}$$

By using the enforced ratio of monomers in phase I and II, see Eq. (S20), we find

$$\frac{\mathrm{d}c_1(t)}{\mathrm{d}t} = \frac{\mathrm{d}c_1^{(\mathrm{I})}(t)}{\mathrm{d}t} \frac{V^{(\mathrm{I})} + \frac{1}{P} V^{(\mathrm{II})}}{V^{(\mathrm{I})}+V^{(\mathrm{II})}}. \tag{S23}$$

Additionally, combining Eq. (S22) with the constitutive equations, Eq. (S16a) and Eq. (S16b), and assuming that $c_\mathrm{a}^{(\mathrm{II})}(t) \approx 0$, we can also write

$$\frac{\mathrm{d}c_1(t)}{\mathrm{d}t} = -2k_+ c_1^{(\mathrm{I})} c_\mathrm{a}^{(\mathrm{I})} \frac{V^{(\mathrm{I})}}{V^{(\mathrm{I})}+V^{(\mathrm{II})}}. \tag{S24}$$

Combining Eq. (S23) and Eq. (S24) finally leads to

$$\frac{\mathrm{d}c_1^{(\mathrm{I})}(t)}{\mathrm{d}t} = -2k_+ c_1^{(\mathrm{I})} c_\mathrm{a}^{(\mathrm{I})} \frac{1}{1 + \frac{1}{P}\frac{V^{(\mathrm{II})}}{V^{(\mathrm{I})}}}. \tag{S25}$$

The aggregate number concentration in phase I follows from Eq. (S16e), given by

$$\frac{\mathrm{d}c_\mathrm{a}^{(\mathrm{I})}(t)}{\mathrm{d}t} = k_1 {c_1^{(\mathrm{I})}}^{n_1} + k_2 {c_1^{(\mathrm{I})}}^{n_2} \left(1 + \frac{1}{P}\frac{V^{(\mathrm{II})}}{V^{(\mathrm{I})}}\right) \left(c_1^{(\mathrm{I})}(0) - c_1^{(\mathrm{I})}(t)\right) \tag{S26}$$

where we have used $M_{\mathrm{a}}^{(\mathrm{II})}(t) \approx 0$ and the mass conservation (see Eq. (S19))

$$\begin{aligned} M_{\mathrm{a}}^{(\mathrm{I})}(t) &= \frac{c_1^{(\mathrm{I})}(0)V^{(\mathrm{I})} + c_1^{(\mathrm{II})}(0)V^{(\mathrm{II})} - c_1^{(\mathrm{I})}(t)V^{(\mathrm{I})} - c_1^{(\mathrm{II})}(t)V^{(\mathrm{II})}}{V^{(\mathrm{I})}} \\ &= \left(1 + \frac{1}{P}\frac{V^{(\mathrm{II})}}{V^{(\mathrm{I})}}\right)\left(c_1^{(\mathrm{I})}(0) - c_1^{(\mathrm{I})}(t)\right). \end{aligned} \tag{S27}$$

The equations (S25) and (S26) have the same form as the homogeneous case introduced in Section S1.1 with the effective rate constants

$$\tilde{k}_1 = k_1, \tag{S28a}$$

$$\tilde{k}_2 = k_2\left(1 + \frac{1}{P}\frac{V^{(\mathrm{II})}}{V^{(\mathrm{I})}}\right), \tag{S28b}$$

$$\tilde{k}_+ = k_+\frac{1}{1 + \frac{1}{P}\frac{V^{(\mathrm{II})}}{V^{(\mathrm{I})}}}. \tag{S28c}$$

The monomer concentrations are then given by

$$c_1^{(\mathrm{I})}(t) = c_1^{(\mathrm{I})}(0)\left[1 + \omega\exp(\kappa t)\right]^{-\theta}, \tag{S29a}$$

$$c_1^{(\mathrm{II})}(t) = \frac{1}{P}c_1^{(\mathrm{I})}(0)\left[1 + \omega\exp(\kappa t)\right]^{-\theta} \tag{S29b}$$

with the parameters

$$\lambda = \sqrt{2\tilde{k}_+\tilde{k}_1 c_1^{(\mathrm{I})}(0)^{n_1}}, \tag{S30a}$$

$$\kappa = \sqrt{2\tilde{k}_+\tilde{k}_2 c_1^{(\mathrm{I})}(0)^{n_2+1}}, \tag{S30b}$$

$$\theta = \sqrt{\frac{2}{n_2(n_2+1)}}, \tag{S30c}$$

$$\omega = \frac{\lambda^2}{2\kappa^2\theta}. \tag{S30d}$$

The aggregate number concentration in phase I then results from Eq. (S16a) and Eq. (S29a) and is given by

$$c_{\mathrm{a}}^{(\mathrm{I})}(t) = \frac{1}{\omega + \exp(-\kappa t)}\frac{\theta\omega\kappa}{2\tilde{k}_+} \tag{S31}$$

and

$$\lim_{t\to\infty} c_{\mathrm{a}}^{(\mathrm{I})}(t) = \frac{\theta\kappa}{2\tilde{k}_+}. \tag{S32}$$

The total aggregate number concentration $c_{\mathrm{a}}(t)$ in the system is given by

$$\begin{aligned} c_{\mathrm{a}}(t) &= \frac{V^{(\mathrm{I})} c_{\mathrm{a}}^{(\mathrm{I})}(t) + V^{(\mathrm{II})} c_{\mathrm{a}}^{(\mathrm{II})}(t)}{V^{(\mathrm{I})} + V^{(\mathrm{II})}} \\ &\approx \frac{V^{(\mathrm{I})}}{V^{(\mathrm{I})} + V^{(\mathrm{II})}} c_{\mathrm{a}}^{(\mathrm{I})}(t) \\ &= \frac{V^{(\mathrm{I})}}{V^{(\mathrm{I})} + V^{(\mathrm{II})}} \frac{1}{\omega + \exp(-\kappa t)} \frac{\theta\omega\kappa}{2\tilde{k}_+}, \end{aligned} \tag{S33}$$

and

$$\lim_{t\to\infty} c_{\mathrm{a}}(t) = \frac{V^{(\mathrm{I})}}{V^{(\mathrm{I})} + V^{(\mathrm{II})}} \frac{\theta\kappa}{2\tilde{k}_+}. \tag{S34}$$

The analytical predictions are compared with numerical solutions of the master equation (see Eq. (1a) and Eq. (1b)) in Figure S3 *D* and shows excellent agreement.

**Aggregation time** Using Eq. (S21), Eq. (S29a) and Eq. (S29b), we derive the time

$$T_{\mathrm{p}} = \frac{1}{\kappa} \ln\left( \frac{(1+\omega)(1-p)^{-\frac{1}{\theta}} - 1}{\omega} \right) \tag{S35}$$

to assemble the fraction $p$ of monomers into linear aggregates.

**Moving front positions** The front position of the moving front in phases I and II are derived analogously to the case of a single phase (see section S1.1) with

$$\eta^{(\mathrm{I})}(t) = \int_0^t \mathrm{d}t' \; 2k_+ c_1^{(\mathrm{I})}(t'), \tag{S36a}$$

$$\eta^{(\mathrm{II})}(t) = \int_0^t \mathrm{d}t' \; 2k_+ c_1^{(\mathrm{II})}(t'), \tag{S36b}$$

and the solution for $\theta = 0.5$ is given by

$$\begin{aligned}
\eta^{(\mathrm{I})}(t) &= \frac{4\tilde{k}_+ c_1^{(\mathrm{I})}(0)}{\kappa} \Big[\tanh^{-1}\left(\sqrt{1+\omega}\right) \\
&\quad - \tanh^{-1}\left(\sqrt{1+\omega \exp(\kappa t)}\right)\Big], && \text{(S37a)} \\
\eta^{(\mathrm{II})}(t) &= \frac{4\tilde{k}_+ c_1^{(\mathrm{I})}(0)}{P\kappa} \Big[\tanh^{-1}\left(\sqrt{1+\omega}\right) \\
&\quad - \tanh^{-1}\left(\sqrt{1+\omega \exp(\kappa t)}\right)\Big], && \text{(S37b)} \\
\eta^{(\mathrm{I})}(\infty) &= \frac{4\tilde{k}_+ c_1^{(\mathrm{I})}(0)}{\kappa} \sinh^{-1}\left(\sqrt{\frac{1}{\omega}}\right), && \text{(S37c)} \\
\eta^{(\mathrm{II})}(\infty) &= \frac{4\tilde{k}_+ c_1^{(\mathrm{I})}(0)}{P\kappa} \sinh^{-1}\left(\sqrt{\frac{1}{\omega}}\right). && \text{(S37d)}
\end{aligned}$$

In Figure S3 *G* we solved the master equation numerically for infinitely high mobilities and show the aggregate concentration profiles as a function of time. The front is clearly visible for both phases and the temporal evolution of its position is in agreement with the analytical predictions, as shown in Fig. S3 *J*.

###### S1.2.3 Rapid transport of monomers and linear aggregates between phases I and II

**Assumptions** We now assume that not only the monomers are rapidly exchanged between the phase with the partitioning coefficient $P$, but also the linear aggregates with $P_\mathrm{f}$, where $P_\mathrm{f}$ is independent of the aggregate size (see Fig. S3 *B*). Then,

$$\begin{aligned}
c_1^{(\mathrm{I})}(t) &= P c_1^{(\mathrm{II})}(t), && \text{(S38a)} \\
c_\mathrm{A}^{(\mathrm{I})}(t) &= P_\mathrm{f} c_\mathrm{A}^{(\mathrm{II})}(t), && \text{(S38b)} \\
M_\mathrm{A}^{(\mathrm{I})}(t) &= P_\mathrm{f} M_\mathrm{A}^{(\mathrm{II})}(t). && \text{(S38c)}
\end{aligned}$$

Additionally we assume that initially, no aggregates are present in the system, corresponding to $M_\mathrm{a}^{(\alpha)}(t=0) = 0$ and $c_\mathrm{a}^{(\alpha)}(t=0) = 0$ for $\alpha = \mathrm{I}, \mathrm{II}$.

**Analytical solutions for the monomer and aggregate number concentrations** Analogously to the previous section, we consider the total monomer concentration $c_1$ (see Eq. (S21) and Eq. (S22)), leading to

$$\frac{\mathrm{d}c_1(t)}{\mathrm{d}t} = \frac{\mathrm{d}c_1^{(\mathrm{I})}(t)}{\mathrm{d}t} \frac{V^{(\mathrm{I})} + \frac{1}{P} V^{(\mathrm{II})}}{V^{(\mathrm{I})} + V^{(\mathrm{II})}}. \tag{S39}$$

Additionally, combining

$$\frac{\mathrm{d}c_1(t)}{\mathrm{d}t} = \frac{\mathrm{d}c_1^{(\mathrm{I})}(t)}{\mathrm{d}t} \frac{V^{(\mathrm{I})}}{V^{(\mathrm{I})} + V^{(\mathrm{II})}} + \frac{\mathrm{d}c_1^{(\mathrm{II})}(t)}{\mathrm{d}t} \frac{V^{(\mathrm{II})}}{V^{(\mathrm{I})} + V^{(\mathrm{II})}} \tag{S40}$$

with Eq. (S16a) and Eq. (S16b) results in

$$\frac{\mathrm{d}c_1(t)}{\mathrm{d}t} = -2k_+ c_1^{(\mathrm{I})} c_\mathrm{a}^{(\mathrm{I})} \frac{V^{(\mathrm{I})} + \frac{1}{P}\frac{1}{P_\mathrm{f}} V^{(\mathrm{II})}}{V^{(\mathrm{I})} + V^{(\mathrm{II})}}. \tag{S41}$$

By comparing Eq. (S39) and Eq. (S41), we derive

$$\frac{\mathrm{d}c_1^{(\mathrm{I})}(t)}{\mathrm{d}t} = -2k_+ c_1^{(\mathrm{I})} c_\mathrm{a}^{(\mathrm{I})} \frac{V^{(\mathrm{I})} + \frac{1}{P}\frac{1}{P_\mathrm{f}} V^{(\mathrm{II})}}{V^{(\mathrm{I})} + \frac{1}{P} V^{(\mathrm{II})}}. \tag{S42}$$

By considering the total aggregate number concentration

$$c_\mathrm{a}(t) = \frac{V^{(\mathrm{I})}}{V^{(\mathrm{I})} + V^{(\mathrm{II})}} c_\mathrm{a}^{(\mathrm{I})}(t) + \frac{V^{(\mathrm{II})}}{V^{(\mathrm{I})} + V^{(\mathrm{II})}} c_\mathrm{a}^{(\mathrm{II})}(t), \tag{S43}$$

$$\frac{\mathrm{d}c_\mathrm{a}(t)}{\mathrm{d}t} = \frac{V^{(\mathrm{I})}}{V^{(\mathrm{I})} + V^{(\mathrm{II})}} \frac{\mathrm{d}c_\mathrm{a}^{(\mathrm{I})}(t)}{\mathrm{d}t} + \frac{V^{(\mathrm{II})}}{V^{(\mathrm{I})} + V^{(\mathrm{II})}} \frac{\mathrm{d}c_\mathrm{a}^{(\mathrm{II})}(t)}{\mathrm{d}t}, \tag{S44}$$

which, by using Eq (S38b), leads to

$$\frac{\mathrm{d}c_\mathrm{a}(t)}{\mathrm{d}t} = \frac{\mathrm{d}c_\mathrm{a}^{(\mathrm{I})}(t)}{\mathrm{d}t} \frac{V^{(\mathrm{I})} + \frac{1}{P_\mathrm{f}} V^{(\mathrm{II})}}{V^{(\mathrm{I})} + V^{(\mathrm{II})}}. \tag{S45}$$

Combining Eq. (S44), Eq. (S16e) and Eq. (S16f), we get

$$\frac{\mathrm{d}c_\mathrm{a}(t)}{\mathrm{d}t} = k_1 {c_1^{(\mathrm{I})}}^{n_1} \frac{V^{(\mathrm{I})} + \frac{1}{P^{n_1}} V^{(\mathrm{II})}}{V^{(\mathrm{I})} + V^{(\mathrm{II})}} + k_2 {c_1^{(\mathrm{I})}}^{n_2} M_\mathrm{a}^{(\mathrm{I})} \frac{V^{(\mathrm{I})} + \frac{1}{P^{n_2}}\frac{1}{P_\mathrm{f}} V^{(\mathrm{II})}}{V^{(\mathrm{I})} + V^{(\mathrm{II})}}. \tag{S46}$$

Since the mass is conserved, we get

$$V^{(\mathrm{I})} \left( c_1^{(\mathrm{I})}(t) + M_\mathrm{a}^{(\mathrm{I})}(t) \right) + V^{(\mathrm{II})} \left( c_1^{(\mathrm{II})}(t) + M_\mathrm{a}^{(\mathrm{II})}(t) \right) = \mathrm{const.} \tag{S47}$$

and if $M_\mathrm{a}^{(\mathrm{I})}(0)=0$ and $M_\mathrm{a}^{(\mathrm{II})}(0)=0$, this results in

$$M_\mathrm{a}^{(\mathrm{I})}(t)=\frac{V^{(\mathrm{I})}+\frac{1}{P}V^{(\mathrm{II})}}{V^{(\mathrm{I})}+\frac{1}{P_\mathrm{f}}V^{(\mathrm{II})}}\left(c_1^{(\mathrm{I})}(0)-c_1^{(\mathrm{I})}(t)\right). \tag{S48}$$

From Eq. (S48), Eq. (S45) and Eq. (S46) we then get

$$\begin{aligned}\frac{\mathrm{d}c_\mathrm{a}^{(\mathrm{I})}(t)}{\mathrm{d}t} &= k_1 {c_1^{(\mathrm{I})}}^{n_1}\frac{V^{(\mathrm{I})}+\frac{1}{P^{n_1}}V^{(\mathrm{II})}}{V^{(\mathrm{I})}+\frac{1}{P_\mathrm{f}}V^{(\mathrm{II})}}+k_2 {c_1^{(\mathrm{I})}}^{n_1}\left[c_1^{(\mathrm{I})}(0)-c_1^{(\mathrm{I})}(t)\right]\\ &\quad\times\frac{V^{(\mathrm{I})}+\frac{1}{P^{n_2}}\frac{1}{P_\mathrm{f}}V^{(\mathrm{II})}}{V^{(\mathrm{I})}+\frac{1}{P_\mathrm{f}}V^{(\mathrm{I})}}\frac{V^{(\mathrm{I})}+\frac{1}{P}V^{(\mathrm{II})}}{V^{(\mathrm{I})}+\frac{1}{P_\mathrm{f}}V^{(\mathrm{II})}}.\end{aligned} \tag{S49}$$

Again, we use the fact that the equations (S42) and (S49) have the same form as in the single phase case introduced in Section S1.1 and can define the effective rate constants

$$\tilde{k}_1 = k_1\frac{V^{(\mathrm{I})}+\frac{1}{P^{n_1}}V^{(\mathrm{II})}}{V^{(\mathrm{I})}+\frac{1}{P_\mathrm{f}}V^{(\mathrm{II})}}, \tag{S50a}$$

$$\tilde{k}_2 = k_2\frac{V^{(\mathrm{I})}+\frac{1}{P^{n_2}}\frac{1}{P_\mathrm{f}}V^{(\mathrm{II})}}{V^{(\mathrm{I})}+\frac{1}{P_\mathrm{f}}V^{(\mathrm{I})}}\frac{V^{(\mathrm{I})}+\frac{1}{P}V^{(\mathrm{II})}}{V^{(\mathrm{I})}+\frac{1}{P_\mathrm{f}}V^{(\mathrm{II})}}, \tag{S50b}$$

$$\tilde{k}_+ = k_+\frac{V^{(\mathrm{I})}+\frac{1}{P}\frac{1}{P_\mathrm{f}}V^{(\mathrm{II})}}{V^{(\mathrm{I})}+\frac{1}{P}V^{(\mathrm{II})}}. \tag{S50c}$$

The solution is given by

$$c_1^{(\mathrm{I})}(t) = c_1^{(\mathrm{I})}(0)\left[1+\omega\exp\left(\kappa t\right)\right]^{-\theta}, \tag{S51a}$$

$$c_1^{(\mathrm{II})}(t) = \frac{1}{P}c_1^{(\mathrm{I})}(0)\left[1+\omega\exp\left(\kappa t\right)\right]^{-\theta}. \tag{S51b}$$

with the parameters

$$\lambda = \sqrt{2\tilde{k}_+\tilde{k}_1\left(c_1^{(\mathrm{I})}(0)\right)^{n_1}}, \tag{S52a}$$

$$\kappa = \sqrt{2\tilde{k}_+\tilde{k}_2\left(c_1^{(\mathrm{I})}(0)\right)^{n_2+1}}, \tag{S52b}$$

$$\theta = \sqrt{\frac{2}{n_2\left(n_2+1\right)}}, \tag{S52c}$$

$$\omega = \frac{\lambda^2}{2\kappa^2\theta}. \tag{S52d}$$

The aggregate number concentration inside and outside of the condensate results from Eq. (S16a) and Eq. (S51a) and is then given by

$$c_{\mathrm{a}}^{(\mathrm{I})}(t) = \frac{1}{\omega + \exp(-\kappa t)} \frac{\theta \omega \kappa}{2\tilde{k}_{+}}, \tag{S53a}$$

$$c_{\mathrm{a}}^{(\mathrm{II})}(t) = \frac{1}{P_{\mathrm{f}}} \frac{1}{\omega + \exp(-\kappa t)} \frac{\theta \omega \kappa}{2\tilde{k}_{+}} \tag{S53b}$$

converging to

$$\lim_{t\to\infty} c_{\mathrm{a}}^{(\mathrm{I})}(t) = \frac{\theta \kappa}{2\tilde{k}_{+}} \tag{S54a}$$

$$\lim_{t\to\infty} c_{\mathrm{a}}^{(\mathrm{II})}(t), = \frac{1}{P_{\mathrm{f}}} \frac{\theta \kappa}{2\tilde{k}_{+}}. \tag{S54b}$$

The total aggregate number concentration in the system is then given by

$$c_{\mathrm{a}}(t) = \frac{V^{(\mathrm{I})} + \frac{1}{P_{\mathrm{f}}} V^{(\mathrm{II})}}{V^{(\mathrm{I})} + V^{(\mathrm{II})}} \frac{1}{\omega + \exp(-\kappa t)} \frac{\theta \omega \kappa}{2\tilde{k}_{+}} \tag{S55}$$

and

$$\lim_{t\to\infty} c_{\mathrm{a}}(t) = \frac{V^{(\mathrm{I})} + \frac{1}{P_{\mathrm{f}}} V^{(\mathrm{II})}}{V^{(\mathrm{I})} + V^{(\mathrm{II})}} \frac{\theta \kappa}{2\tilde{k}_{+}}. \tag{S56}$$

In Fig. S3 *E* we compare our analytical predictions for the monomer concentration and the aggregate number concentration with the numerical solution of the master equation for infinitely high mobilities and we find that both are in excellent agreement.

**Aggregation time** It takes the time

$$T_{\mathrm{p}} = \frac{1}{\kappa} \ln\left( \frac{(1+\omega)(1-p)^{-\frac{1}{\theta}} - 1}{\omega} \right) \tag{S57}$$

to assemble a fraction $p$ of all monomers.

**Moving front positions** The front position of the moving front in phase I is derived analogously to the case of a single phase (see section S1.1) with

$$\eta^{(\mathrm{I})}(t) = \int_0^t \mathrm{d}t'\ 2\tilde{k}_{+} c_1^{(\mathrm{I})}(t'). \tag{S58}$$

Note that here we use $\tilde{k}_+$ instead of $k_+$ since the aggregate concentrations in both phases are coupled (see Eq. (S38b)). The solution for $\theta = 0.5$ is given by

$$\eta^{(\mathrm{I})}(t) = \frac{4\tilde{k}_+ c_1^{(\mathrm{I})}(0)}{\kappa}\Big[\tanh^{-1}\left(\sqrt{1+\omega}\right) - \tanh^{-1}\left(\sqrt{1+\omega\exp\left(\kappa t\right)}\right)\Big], \tag{S59}$$

$$\eta^{(\mathrm{I})}(\infty) = \frac{4\tilde{k}_+ c_1^{(\mathrm{I})}(0)}{\kappa}\sinh^{-1}\left(\sqrt{\frac{1}{\omega}}\right). \tag{S60}$$

Since for $P > 1$ and $P_\mathrm{f} > 1$, in phase I we expect more material, the front position in this phase will also be larger. Since we allow for rapid material exchange between the inside and outside of the condensate, the front will be located at the same location in phase II as in phase I:

$$\eta^{(\mathrm{II})}(t) = \eta^{(\mathrm{I})}(t). \tag{S61}$$

This result can be confirmed by numerically solving the master equation and investigating the aggregate concentration profile (see Fig. S3 *H* and Fig. S3 *K*) and indeed, the front positions are identical for both phases.

**Behaviour in the limit $P_\mathrm{f} \to \infty$** For large aggregate partitioning coefficient $P_\mathrm{f} \to \infty$ (see Fig. S3 *C*), the effective rate constants are

$$\tilde{k}_1 = k_1\left(1 + \frac{1}{P^{n_1}}\frac{V^{(\mathrm{II})}}{V^{(\mathrm{I})}}\right), \tag{S62a}$$

$$\tilde{k}_2 = k_2\left(1 + \frac{1}{P}\frac{V^{(\mathrm{II})}}{V^{(\mathrm{I})}}\right), \tag{S62b}$$

$$\tilde{k}_+ = k_+\frac{1}{1 + \frac{1}{P}\frac{V^{(\mathrm{II})}}{V^{(\mathrm{I})}}}. \tag{S62c}$$

The rate constants $\tilde{k}_2$ and $\tilde{k}_+$ are identical to the case without aggregate exchange between the phases, see Eq. (S28b) and Eq. (S28c). Thus, the characteristic time of aggregation $\kappa$ is identical for these two cases. With these rate constants, the monomer concentration and aggregate number concentration is in agreement with the analytical predictions for general $P_\mathrm{f}$ (see Fig. S3 *F*).

The front position results from

$$\eta^{(\mathrm{I})}(t) = \int_0^t \mathrm{d}t'\ 2k_+ c_1^{(\mathrm{I})}(t'). \tag{S63}$$

Note that for the calculation of $\kappa$ and $\lambda$ we use $\tilde{k}_+$, but in the front position integral we use $k_+$. This results from the fact that the only location where aggregates exist and thus, linear aggregate elongation takes place, is located in phase I and we can treat it as a single phase (see Appendix S1.1). For $\theta = 0.5$, the front is then given by

$$\begin{aligned}\eta^{(\mathrm{I})}(t) &= \frac{4k_+ c_1^{(\mathrm{I})}(0)}{\kappa}\Big[\tanh^{-1}\left(\sqrt{1+\omega}\right) \\ &\quad - \tanh^{-1}\left(\sqrt{1+\omega\exp(\kappa t)}\right)\Big], \qquad \text{(S64)} \\ \eta^{(\mathrm{I})}(\infty) &= \frac{4k_+ c_1^{(\mathrm{I})}(0)}{\kappa}\sinh^{-1}\left(\sqrt{\frac{1}{\omega}}\right). \qquad \text{(S65)}\end{aligned}$$

In phase II we assume that for $P_{\mathrm{f}} \to \infty$ no aggregates can be found and thus, no front exists. This can be confirmed with numerical solutions of the master equation (see Fig. S3 *I* and Fig. S3 *L*).
